# Drosophila Models Uncover Substrate Channeling Effects on Phospholipids and Sphingolipids in Peroxisomal Biogenesis Disorders

**DOI:** 10.1101/2024.04.26.591192

**Authors:** Michael F Wangler, Yu-Hsin Chao, Mary Roth, Ruth Welti, James A. McNew

**Affiliations:** Molecular and Human Genetics, Baylor College of Medicine, Houston TX; Jan and Dan Duncan Neurological Research Institute, Texas Children’s Hospital; Kansas Lipidomics Research Center, Division of Biology, Kansas State University, Manhattan, KS; Department of BIoSciences, Rice University

**Keywords:** Peroxisome, phospholipid, sphingolipid, *Drosophila*

## Abstract

Peroxisomal Biogenesis Disorders Zellweger Spectrum (PBD-ZSD) disorders are a group of autosomal recessive defects in peroxisome formation that produce a multi-systemic disease presenting at birth or in childhood. Well documented clinical biomarkers such as elevated very long chain fatty acids (VLCFA) are key biochemical diagnostic findings in these conditions. Additional, secondary biochemical alterations such as elevated very long chain lysophosphatidylcholines are allowing newborn screening for peroxisomal disease. In addition, a more widespread impact on metabolism and lipids is increasingly being documented by metabolomic and lipidomic studies. Here we utilize *Drosophila* models of *pex2* and *pex16* as well as human plasma from individuals with *PEX1* mutations. We identify phospholipid abnormalities in *Drosophila* larvae and brain characterized by differences in the quantities of phosphatidylcholine (PC) and phosphatidylethanolamines (PE) with long chain lengths and reduced levels of intermediate chain lengths. For diacylglycerol (DAG) the precursor of PE and PC through the Kennedy pathway, the intermediate chain lengths are increased suggesting an imbalance between DAGs and PE and PC that suggests the two acyl chain pools are not in equilibrium. Altered acyl chain lengths are also observed in PE ceramides in the fly models. Interestingly, plasma from human subjects exhibit phospholipid alterations similar to the fly model. Moreover, human plasma shows reduced levels of sphingomyelin with 18 and 22 carbon lengths but normal levels of C24. Our results suggest that peroxisomal biogenesis defects alter shuttling of the acyl chains of multiple phospholipid and ceramide lipid classes, whereas DAG species with intermediate fatty acids are more abundant. These data suggest an imbalance between *de novo* synthesis of PC and PE through the Kennedy pathway and remodeling of existing PC and PE through the Lands cycle. This imbalance is likely due to overabundance of very long and long acyl chains in PBD and a subsequent imbalance due to substrate channeling effects. Given the fundamental role of phospholipid and sphingolipids in nervous system functions, these observations suggest PBD-ZSD are diseases characterized by widespread cell membrane lipid abnormalities.

## Introduction

Peroxisomal biogenesis is an evolutionarily conserved process that allows the cell to generate functional peroxisomes. Genetic mutations in the peroxisomal biogenesis machinery, affecting proteins which are encoded by the *PEX* genes in humans, lead to global biochemical defects that reflect lack of peroxisomes (Wanders and Waterham 2005; Braverman et al. 2016). The lack of functional peroxisomes leads to a number of metabolic changes which relate directly to peroxisomal biochemistry such as elevations in very long chain fatty acids (VLCFAs), reduced plasmalogens, and increased phytanic and pipecolic acid (Wanders and Waterham 2005; Moser et al. 1999). Recently, newborn screening for peroxisomal disease, particularly X-linked Adrenoleukodystrophy has been initiated in multiple countries based on secondary biochemical alterations in peroxisomal dysfunction impacting lysophosphatidylcholines (Braverman et al. 2016; Mallack et al. 2021; Klouwer et al. 2017). This screening ultimately depends on an imbalance in the length of acyl chains observed when peroxisomal fatty acid metabolism is altered or absent, stemming from the defect in peroxisomal beta-oxidation and leading to an over-abundance of very long acyl chains.

Metabolomic and lipidomic studies on peroxisomal disorders have documented the metabolic consequences of general peroxisomal defects and uncovered additional abnormalities in peroxisomal disorders beyond the classical peroxisomal biochemical markers (Herzog et al. 2017, 2016; Wangler et al. 2018, 2023). These studies have demonstrated that a generalized defect in peroxisomal biochemistry is not restricted to affecting peroxisomal pathways but also produces a myriad of downstream secondary biochemical changes. Of note, the phospholipid compositions of tissues from patients with PBD-ZSD are altered and display increased chain length and increased number of unsaturations in PBD (Herzog et al. 2016). Indeed, altered phospholipids have been shown to correlate with other biomarkers of peroxisomal disease (Herzog et al. 2018). This has also affected sphingomyelin levels in fibroblasts of PBD-ZSD versus controls (Herzog et al. 2016). In our previous metabolomic studies we also observed reduced plasma sphingomyelin levels in PBD-ZSD (Wangler et al. 2018).

Phosphatidylethanolamine (PE) and phosphatidylcholine (PC) are the most abundant phospholipids and their synthesis is mediated in part by the Kennedy pathway which allows *de novo* synthesis of phospholipids from diacylglycerols (DAG) (Gibellini and Smith 2010). The biochemical steps for PE and PC synthesis do not occur within peroxisomes and so the effects on these lipids are examples of secondary alterations in metabolites that are not primarily peroxisomal in etiology. When considering these secondary metabolic consequences of PBD there are several considerations. Some of the secondary metabolic abnormalities could stem from a primary excess or deficiency of a substrate that comes from peroxisomal metabolism such as excess very long chain fatty acid or deficient plasmalogens. In addition, regulation of overall cellular metabolism could be affected in PBD which could lead to secondary metabolic changes. In order to explore the full extent of the metabolic consequences of PBD one approach is to leverage animal models of these disorders and to perform comprehensive metabolic or lipidomic assessments. We have used *Drosophila* models such as *pex2* and *pex16* as well as *pex3* to fully catalogue the impact of peroxisomal biogenesis defects on metabolism (Wangler et al. 2017; Faust et al. 2014). Here we extend these analyses using a thorough phospholipid analysis (Spivey et al. 2023; Lusk et al. 2022) including PC and PE and their precursor DAGs in addition to ceramides and PE ceramides and sphingomyelin.

## Materials and Methods

### Fly husbandry

All flies were maintained at room temperature (21°C) and except where otherwise noted experiments were conducted at room temperature. The *pex2^1^*, and *pex2^2^* lines were derived from imprecise excision of (*w[1118]*; P{w[+mC] = EPg}*pex2*[HP35039]/TM3, Sb[1], see below) these were then backcrossed 5 generations with *y w*: FRT80B and studied as: *y w*; FRT80B *y w*; FRT80B-*pex2^1^* y w; FRT80B-*pex2^2^w[1118*];PBac{y[+mDint2]w[+mC] = 53M21}VK00037;FRT80B-*pex2^2^* w[1118];PBac{y[+mDint2]w[+mC] = 53M21}VK00037;FRT80B-*pex2^1^* Except where otherwise indicated the 5 strains above each crossed to a genomic deficiency uncovering *pex2* locus *w1118*; Df(3L)BSC376/TM6C,*Sb1 cu1* are labeled as *pex2* Control, *pex2^2^, pex2^1^*, *pex2^2^*Rescue, and *pex2^1^* Rescue, respectively.

The *y w*: *pex16^1^* line [41] was obtained from Kenji Matsuno, derived from: *y 1 w67c23*; P{GSV6}*pex16*GS14106/TM3, Sb1 Ser1. The *y w*: *pex16*^EY^ strain was obtained from Bloomington Stock center *y[1] w[67c23]*; P{*w* [+mC] y[+mDint2] = EPgy2}*Pex16*[EY05323]. *y w*; FRT80B y w; pex161 y[1] w[67c23]; P{w[+mC] y[+mDint2] = EPgy2}pex16[EY05323] w[1118];PBac{y[+mDint2]w[+mC] = 115M13}VK00037;*pex16^1^*w[1118];PBac{y[+mDint2]w[+mC] = 115M13}VK00037;P{w[+mC]. Except where otherwise indicated, the 5 strains above each crossed to a genomic deficiency uncovering the *pex16* locus *w1118*; Df(3L)BSC563/TM6C,*cu1 Sb1* are labeled as *pex16* Control, *pex16^1^ pex16^EY^ pex16^1^* Rescue and *pex16^EY^*Rescue respectively.

### Human Samples

All subjects were recruited to an institutional review board–approved study, “The Biochemical and Cell Biology of Peroxisomal Disorders Study,” at Baylor College of Medicine (H-32837) or “Translational Models of Neurological Disease Study” (H-44779)(Wangler et al. 2023; Bacino et al. 2015; Ventura et al. 2016). All subject caregivers/parents provided informed consent, including consent to publish patient photos. All participants were examined, and samples were collected under the same conditions. Ascertainment was based on the presence of molecularly confirmed mutations in the PEX genes and/or biochemical conifrmation of a defect in peroxisomal biogenesis. Blood samples were collected by a trained phlebotomist and immediately plasma was isolated by ultracentrifugation. Each plasma sample was divided into aliquots and frozen at −80 ° C.

### Phospholipid and diacylglycerol analysis

*Drosophila* phospholipids and diacylglycerols were analyzed by direct-infusion into an electrospray ionization triple quadrupole mass spectrometer (Applied Biosystems 4000 Q-trap mass spectrometer, Sciex, Framingham, MA, US), as described (Scheitz et al. 2013). The plasma phospholipids (3 μL plasma per sample) were analyzed similarly using a Water Xevo TQS mass spectrometer (Waters Corp., Milford, MA, US). Preparation of plasma, internal standard information, and details are presented in Zhou et al. (2012). Mass spectrometry parameters on the Xevo TQS were source temperature, 150°C; desolvation temperature, 250°C; cone gas flow, 150 L h^-1^; desolvation gas flow, 650 L h^-1^; collision gas flow, 0.14 mL min^-1^; nebulizer gas, 7 Bar; LM 1 resolution, 3.2; HM 1 resolution, 15.5. Scan and data processing parameters are shown in **Supplemental Table S1**.

## Results

### Drosophila pex2 and pex16 mutant lines

*Drosophila pex2* and *pex16* null mutants have been previously studied as peroxisomal loss-of-function models in flies and we utilized these mutant and genomic rescue controls in this study (Wangler et al. 2017). For *pex16* we utilize a deletion (Nakayama et al. 2011) and an EY insertion P-element line (Bellen et al. 2004). For pex2 both alleles utilized are considered null alleles that delete the gene. In the case of *pex16*, the *pex16* deletion allele is also considered a null, although the EY insertion preserves the coding sequence and has some attenuated phenotypes suggesting it is a hypomorph. In order to study *pex16* (**Figure 1A**) and *pex2* (**Figure 1B**) mutants, we utilize genomic rescue lines and controls as described (Wangler et al. 2017). Because we had previously used untargeted metabolomics and observed phospholipid abnormalities (**Figure S1**), we pursued lipidomic analysis of the *Drosophila pex2* and *pex16* mutants in larvae as well as adult heads (primarily Drosophila adult brain) (**Figure 1C**).

**Figure 1.**
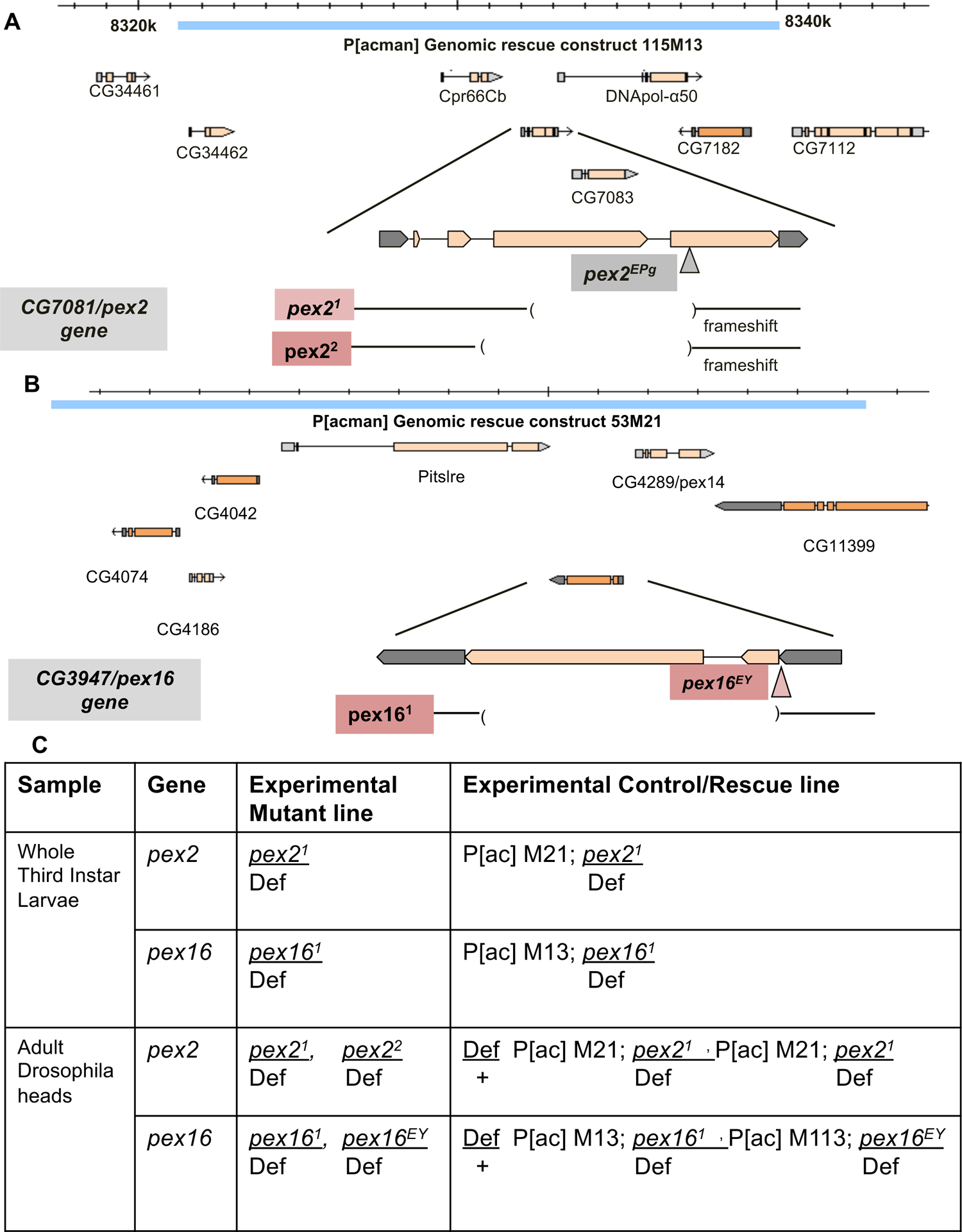
Genotypes of Drosophila peroxisomal biogenesis mutant lines used in the lipidomic study A. The *pex2* gene (former CG7081) in *Drosophila melanogaster* is on 3L and is depicted on the chromosomal region. The *pex2^1^*, and *pex2^2^* lines are deletion alleles. The genomic rescue construct line is derived from 115M13 clone and rescues the entire region (blue bar). B. The *pex16* gene (former CG3947) in *Drosophila melanogaster* is on 3L and is depicted on the chromosomal region. The *y w*: *pe16^1^* line is depicted as a deletion allele ( c/o Kenji Matsuno). The *y w: pex16^EY^* strain is an EY insertion allele in the 5’UTR of the gene. The genomic rescue construct line is derived from the 53M21 clone (blue bar). C. The *Drosophila* experiments analyzed in this study were conducted on whole third instar larvae and then another set of experiments on adult *Drosophila* heads (brain and cuticle). The genotypes for each of these experiments include analysis of the *pex2^1^* and the *pe16^1^* allele were studied in comparison to genomics rescue lines. For the adult head experiments two alleles each for *pex2* and *pex16* were studied in comparison to both control and rescue lines.

### Drosophila pex mutant larvae exhibit altered chain lengths of phosphatidylcholine and phosphatidylethanolamine

First, we examined the *pex* mutant at the *Drosophila* larval stage and focused on phospholipid analysis. We observed that the relative amounts (in mol %) of phosphatidylethanolamine (PE) (**Figure 2A**), and phosphatidylcholine (PC) (**Figure 2B**) either exhibited minor differences or are equivalent between *pex2* mutant and rescue and *pex16* mutant and rescue. This was consistent with our previous study using untargeted metabolomic methods showing no difference in quantity for PC and PE (Wangler et al. 2017). However, while the total of all sub-types appears unchanged in *pex2* and *pex16* mutants, specific subtypes are altered and there is a clear pattern of these alterations related to the carbon chain length. Both PE 30:1 (**Figure 2C**) and PC 30:1 (**Figure 2D**) are reduced in *pex2* mutant compared to rescue and *pex16* mutant compared to rescue, although the difference was not statistically significant for *pex2* for PE 30:1. Note that in this nomenclature for PC and PE, the 30:1 means that the two acyl chains have a total of 30 carbons and 1 unsaturation total. Across the spectrum of phospholipids measured, all the acyl groups are likely between C12 and C22 and would be considered long chain fatty acids or acyl groups. However, we observed a pattern specific to 28 and 30 carbon that differed from 38 carbon, for example. In order to characterize this, we distinguish between broad classes of long chain phospholipids and so we term a phospholipid to have an intermediate chain length if the total carbon chain (across the two acyl chains) is C28-C31, and we consider C32 and above to be long chain length phospholipids. Indeed, a number of intermediate chain length PC and PE’s exhibit the same pattern. PC 28:1 and PC 28:0 are both reduced in *pex2* mutants compared to rescue and *pex16* mutant compared to rescue (**Figure S2A**). PE 28:1 and PE 28:0 also show reduction in the *pex2* and *pex16* mutants compared to rescue larvae (**Figure S2B**). Reduced levels of other intermediate chain phospholipids such as PC 30:2, PC 30:0, PC 31:2, PC 31:1 and PC 31:0 are also seen (**Figure S2C**).

**Figure 2.**
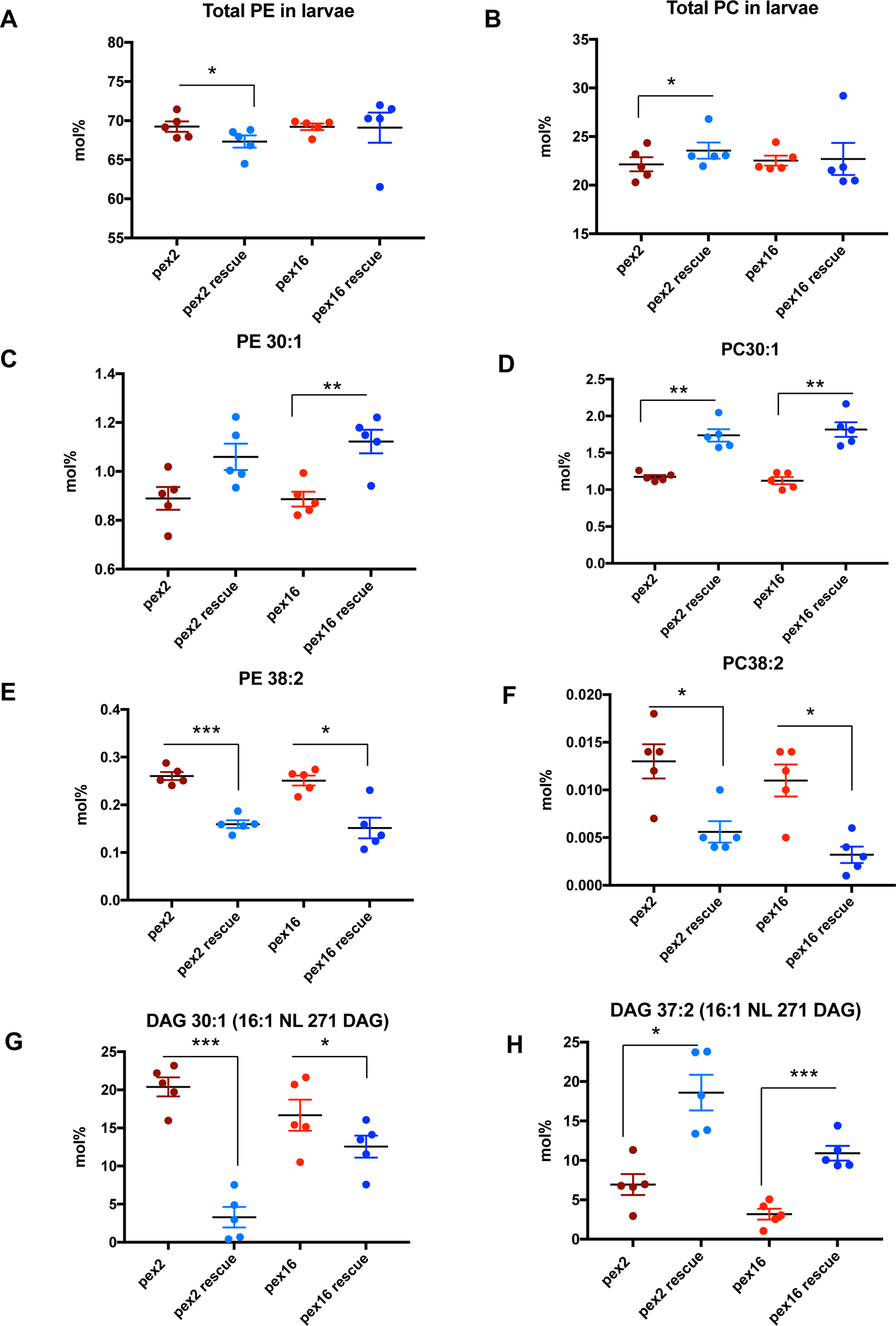
Chain-length specific phospholipid and diacylglycerol abnormalities in peroxisomal mutant larvae. A. Total levels in mol% of phosphatidylethanolamine (PE) in *pex2* and *pex16* larvae shows a mild but statistically significant increase in total PE in *pex2* mutant larvae compared to pex2 rescue (ratio *pex2/pex2 rescue*: 1.03, p=0.0221), and no difference in PE in *pex16* mutant larvae compared to pex16 rescue (pex16/pex16 rescue: ratio 1.00, p=0.94). B. Total levels in mol% of phosphatidylcholine (PC) in *pex2* and *pex16* larvae shows a mild but statistically significant decrease in total PC in *pex2* mutant larvae compared to *pex2* rescue (ration *pex2/pex2* rescue: ratio 0.94,p=0.038), and no difference in PC in *pex16* mutant larvae compared to *pex16* rescue (*pex16/pex16* rescue: ratio 0.99, p=0.897). C. Levels in mol% of PE 30:1 in *pex2* and *pex16* larvae shows no significant difference in PE 30:1 in *pex2* mutant larvae compared to pex2 rescue (ratio *pex2/pex2 rescue*: 0.839, p=0.068), and a significant reduction in *pex16* mutant larvae compared to pex16 rescue (pex16/pex16 rescue: ratio 0.790, p=0.008). D. Levels in mol% of PC 30:1 in *pex2* and *pex16* larvae shows a significant decrease in PC 30:1 in *pex2* mutant larvae compared to pex2 rescue (ratio *pex2/pex2 rescue*: 0.675, p=0.0005), and a significant reduction in in *pex16* mutant larvae compared to *pex16* rescue (pex16/pex16 rescue: ratio 0.618, p=0.002). E. Levels in mol% of PE 38:2 in *pex2* and *pex16* larvae shows a significant increase in PE 38:2 in *pex2* mutant larvae compared to pex2 rescue (ratio *pex2/pex2 rescue*: 1.63, p=0.0003), and a significant reduction in in *pex16* mutant larvae compared to *pex16* rescue (pex16/pex16 rescue: ratio 1.660, p=0.011). F. Levels in mol% of PC 38:2 in *pex2* and *pex16* larvae shows a significant increase in PC 38:2 in *pex2* mutant larvae compared to pex2 rescue (ratio *pex2/pex2 rescue*: 2.303, p=0.043), and a significant reduction in in *pex16* mutant larvae compared to *pex16* rescue (pex16/pex16 rescue: ratio 3.24, p=0.019). G. Levels in mol% of Diacylglycerol 30:1 (16:1 NL 271 DAG) in *pex2* and *pex16* larvae shows a significant increase in DAG 30:1 in *pex2* mutant larvae compared to pex2 rescue (ratio *pex2/pex2 rescue*: 6.197, p=0.0006), and a significant reduction in in *pex16* mutant larvae compared to *pex16* rescue (pex16/pex16 rescue: ratio 1.328, p=0.0477). H. Levels in mol% of Diacylglycerol 37:2 (16:1 NL 271 DAG) in *pex2* and *pex16* larvae shows a significant decrease in DAG 37:2 in *pex2* mutant larvae compared to pex2 rescue (ratio *pex2/pex2 rescue*: 0.373, p=0.0257), and a significant reduction in in *pex16* mutant larvae compared to *pex16* rescue (pex16/pex16 rescue: ratio 0.291, p=0.0003).

In contrast, long chain length PCs and PEs exhibit the opposite effect. Both *pex2* and *pex16* mutants have increased PE 38:2 and PC 38:2 phospholipids (**Figure 2E-F**). Likewise PC 37:2 is also increased in peroxisome deficient larvae (**Figure S3)**. These data suggest that while the overall pool of PCs and PEs in peroxisome deficient mutant larvae appears similar to controls and rescue, there is a deficiency of intermediate chain-length species and excess of long chain lengths. We therefore hypothesized that in peroxisomal mutants due to the excess of very long chain fatty acids, an over-abundance of long acyl chains would result.

One possibility to explain the increased abundance of long chains and reduced intermediate chain lengths is that altered chain lengths of the precursors of PCs and PEs might produce an imbalance in downstream products. We therefore examined diacylglycerols (DAGs) which can be converted to PCs and PEs through the Kennedy pathway (Gibellini and Smith 2010). Counter to our initial expectation, we observed an apparent excess of intermediate chain such as DAG 30:1 in both *pex2* and *pex16* mutants (**Figure 2G**). Long chain length DAGs such as DAG 37:2 were reduced in *pex2* and *pex16* mutants (**Figure 2H**). These data suggest that intermediate chain length phospholipid DAGs are actually increased in the presence of decreased PC and PEs in *pex* mutants. Thus, the DAG precursors exhibit an opposite pattern compared to PE and PC in terms to which chain lengths are most abundant in peroxisomal mutants.

The overall patterns are apparent when looking across multiple lipids, while individual lipids could vary. We plotted the ratio of specific phospholipids for *pex2* mutants compared to *pex2* rescue and *pex16* mutants compared to *pex16* rescue. The ratio of PCs across a range of chain lengths from PC 28:1 to PC 38:1 showed an increase in PC amounts as chain length increases (**Figure 3A**), with similar findings for PEs (**Figure 3B**).

**Figure 3.**
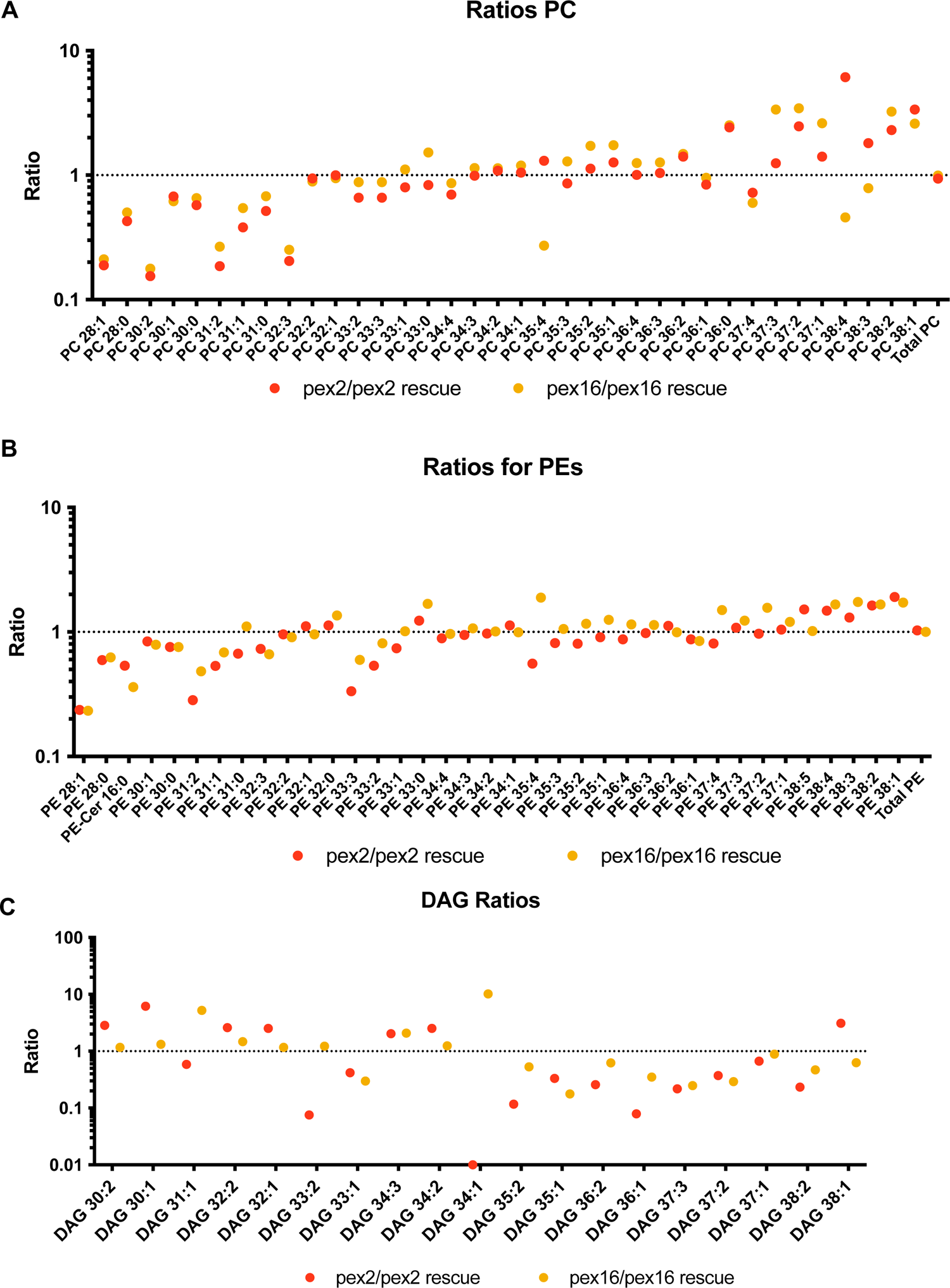
Inverse relationship between phospholipid and diacylglycerol ratios spanning intermediate and long chain lengths. A. Ratios of PCs comparing *pex2/pex2* rescue (red) and *pex16/pex16* rescue (yellow). These ratios are notably low for intermediate chain length PCs. For PC 28:1, pex2/pex2 rescue ratio 0.1889, pex16/pex16 rescue ratio 0.2111. For PC 30:2 *pex2/pex2* rescue ratio 0.154, *pex16/pex16* rescue ratio 0.177. These ratios gradually increase with variability as the chain length increases. For PC 38:2, pex2/pex2 rescue ratio 2.303, pex16/pex16 rescue ratio 3.243. For PC 38:1, pex2/pex2 rescue ratio 3.372, pex16/pex16 rescue 2.598. B. Ratios of PEs comparing pex2/pex2 rescue (red) and pex16/pex16 rescue (yellow). These ratios are notably low for intermediate chain length PEs. For PE 28:1, pex2/pex2 rescue ratio 0.2369, pex16/pex16 rescue ratio 0.2328. For PE 31:2 *pex2/pex2* rescue ratio 0.2839, pex16/pex16 rescue ratio 0.4841. These ratios gradually increase with variability as the chain length increases. For PE 38:2, *pex2/pex2* rescue ratio 1.631, pex16/pex16 rescue ratio 1.660. For PE 38:1, pex2/pex2 rescue ratio 1.906, pex16/pex16 rescue 1.723. C. Ratios of DAGs comparing *pex2/pex2* rescue (red) and *pex16/pex16* rescue (yellow). These ratios are notably high for intermediate chain length (in contrast to PEs and PCs). For DAG 30:1 *pex2/pex2* rescue ratio 6.197, pex16/pex16 rescue ratio 1.328. DAG 32:2 *pex2/pex2* rescue ratio 2.601 pex16/pex16 rescue ratio 1.474. These ratios gradually decrease with variability as the chain length increases (in contrast to the increases seen in PEs and PCs). For DAG 37:2, *pex2/pex2* rescue ratio 0.3726, pex16/pex16 rescue ratio 0.2913. For DAG 38:2, pex2/pex2 rescue ratio 1.906, pex16/pex16 rescue 1.723.

These ratios show that the phospholipids have an imbalance toward long chain lengths in the mutants. In contrast, DAG ratios in *pex* mutants have an imbalance toward intermediate chain lengths (**Figure 3C**). These observations have some similarities to the cellular data of patients with PBD previously reported (Herzog et al. 2016), and in those studies, not only chain-length but also the number of unsaturations in the carbon chain have also been observed to play a role. We therefore examined the phospholipids according to how many unsaturations in *Drosophila* mutants (**Figure S4**). For example examining PC 30:0, PC 32:0 and PC 34:0 as a group of PCs with zero unsaturations (e.g. PC N:0). The ratio of *pex2* and *pex16* mutants compared to rescue show increases as chain length increases across different numbers of unsaturations such as PC N:0, PC N:1, PC N:2 and PC N:3 phospholipid (where N represents the carbon chain of different lengths), but the number of unsaturations themselves did not appear to have a dramatic impact on levels in *Drosophila pex* mutant larvae (**Figure S4**). In summary, the *Drosophila* larvae have PE and PC with longer carbon chain lengths and DAG precursors with more intermediate carbon chain lengths. Moreover, in *Drosophila* larvae the number of unsaturations does not appear to be dramatically altered in the mutants.

### Phospholipid alterations in the brain of Drosophila pex mutant adult flies

Next we examined the adult fly brain to assess the neuro-metabolic impact in *pex2* and *pex16* mutants. In contrast to larvae where the relative amounts of PE and PC were similar between *pex* mutants and rescue lines, altered relative amounts of total PE and PC were observed in adult heads. For *pex2*, both mutant alleles were associated with reduced relative amounts of total PE compared to rescue lines, although the mutants’ PE levels were not significantly different than the control line. This discrepancy could be due to the control line not matching the genetic background, and in general the genomic rescue line could be regarded as a more appropriate control for this background (**Figure 4A**). Similarly for *pex16*, reduced total PE is observed in the *pex16^1^* allele compared to rescue, with a much less dramatic (though still significant) difference between the hypomorph *pex16^EY^* line and rescue (**Figure 4A**). The difference between the two *pex16* mutant lines is likely due to comparison between a null and hypomorphic allele (Wangler et al. 2017). For total PC in the adult brain there were clear differences between all the pex2 alleles with higher levels of PC comopared to control and rescue lines (**Figure 4B**). For pex16 there was a complex range of results for the different genotypes (**Figure 4B**). Taken together these data suggest there may be an overall reduction in total PE in adult *Drosophila* brain, although variability amongst control and rescue lines lead to uncertainty about the biological significance of these alterations.

**Figure 4.**
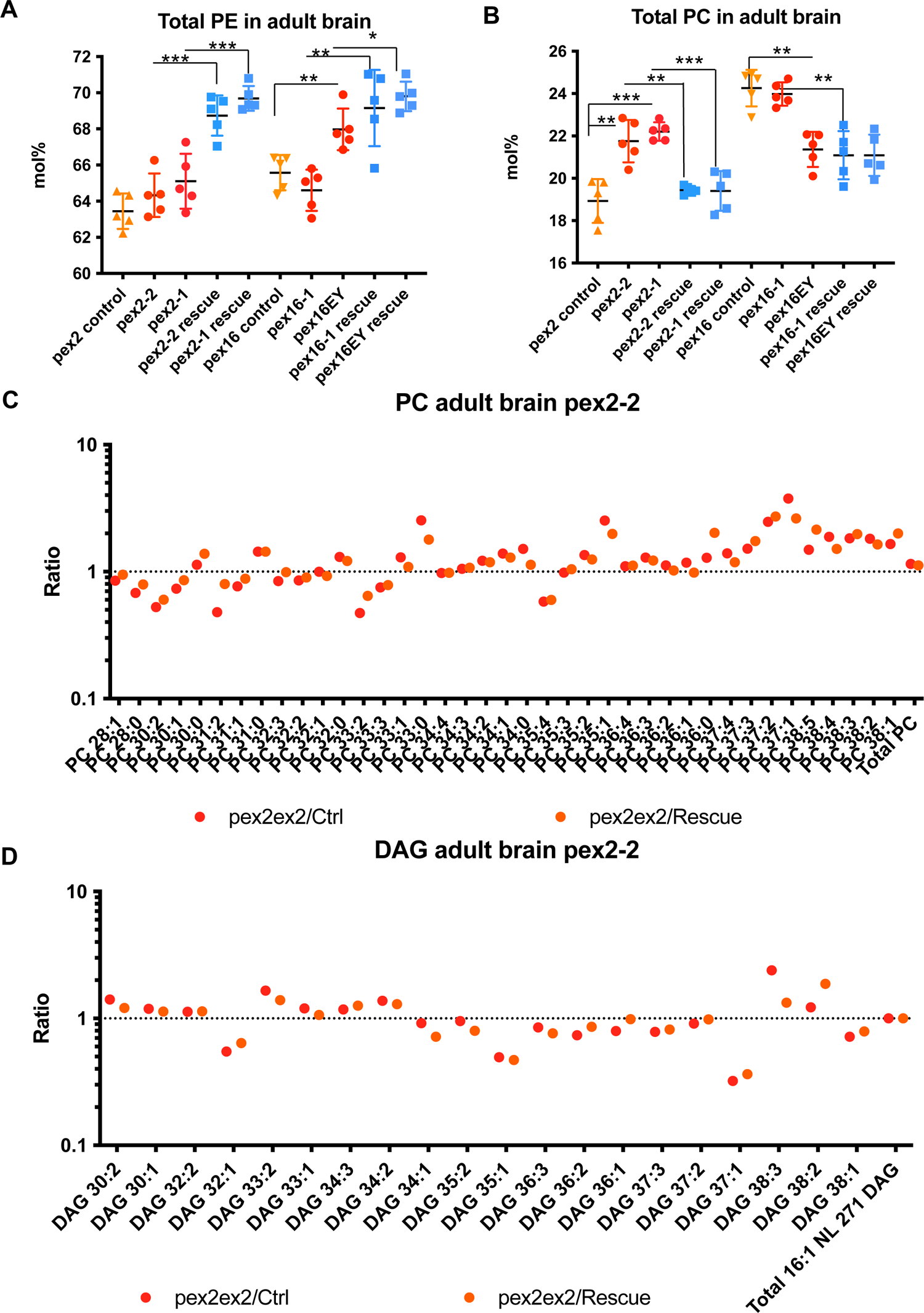
Chain-length specific phospholipid and diacylglycerol abnormalities in peroxisomal mutant brain. A. Total levels in mol% of Phosphatidylethanolamine (PE) in *pex2* and *pex16* adult brains (genotypes shown in Figure 1) shows a reduction in total PE in *pex2* mutant brain compared to pex2 rescue (ratio *pex2^2/^pex2^2^* rescue 0.936, p=<0.0001; ratio *pex2^1^/pex2^1^* rescue 0.934, p=0.001), and reduced PE in *pex16* mutant larvae compared to pex16 rescue (*pex16^1^/pex16^1^* rescue 0.934, p=0.005; ratio pex16EY/Pex16EY rescue 0.974, p=0.0226). B. Total levels in mol% of Phosphatidylcholine (PC) in *pex2* and *pex16* adult brains (genotypes shown in Figure 1) shows increased levels in total PC in pex2 mutant brain compared to pex2 rescue (ratio *pex2^2/^pex2^2^*rescue 1.119, p=0.006; ratio *pex2^1^/pex2^1^* rescue 1.144, p=0.001), but the effect on *pex16* mutant larvae is complex with the significant differences between *pex16^EY^* as being reduced compared to control (but not *pex16^EY^* rescue) and *pex16^1^* levels being increased compared to *pex16^1^* rescue (but not control). C. Ratios of PCs comparing *pex2^2^/Control* (red) and *pex2^2^/pex2^2^* rescue (yellow). These ratios are notably low for some intermediate chain length PCs. These ratios gradually increase with variability as the chain length increases. D. Ratios of DAGs comparing *pex2^2^/Control* (red) and *pex2^2^/pex2^2^* rescue (yellow). These ratios are generally higher for intermediate chain length (in contrast to PEs and PCs). These ratios gradually decrease with variability as the chain length increases (in contrast to the increases seen in PEs and PCs).

We also examined whether the *pex* mutant brains have the same chain length alterations as we observed in the larvae. For each mutant line (*pex2^1^*), (*pex2^2^*), (*pex 16^1^*), (*pex16^EY^*) we compared ratios of PC for mutant and rescue and mutant and control across chain lengths. For example, *pex2^2^*mutants have reduced ratios for intermediate chain lengths compared to both rescue and control lines and increased ratios for longer chain lengths, an effect which is similar to that seen in larvae (**Figure 4C**). While the overall change in ratios is less dramatic, we also observed higher levels of DAGs of intermediate chain length in *pex* mutant brain and lower levels of DAG of longer chain length (**Figure 4D**). The most consistent phospholipid alterations in *Drosophila* brain could be grouped according to PCs that were consistently decreased in peroxisome deficiency mutants such as PC 30:1 and PC 30:2 (**Figure S5**), versus PCs that were consistently increased in peroxisomal mutants such as PC 30:0, PC 35:1, PC 34:1, PC 34:2, PC 37:2 and PC 37:1 (**Figure S6**). PE’s exhibited different patterns with consistent decreases in PE 33:3, PE 33:2, PE 33:1, PE 35:4, PE 35:3 and PE 35:2 (**Figure S7**). Consistent increases in PE were observed in PE 33:0, PE 34:4, PE 34:3, PE 34:1, PE 36:3 and PE 38:3 (**Figure S8**). These findings pointed to phospholipid chains where the number of unsaturations could influence whether the levels were increased or decreased. Nonetheless some similar trends of increasing levels with increasing chain lengths could be observed for pex2 and pex16 null alleles (**Figure S9**). These findings suggest the altered chain length of PEs and PC’s and the reciprocal abnormalities for DAG is not only observed in larvae but can be seen in adult fly brain also.

### Altered PE ceramides in pex mutant brains

In a previous metabolomic analyses in patients with PBD-ZSD, typically with mutations in the *PEX1* gene, we observed reduced sphingomyelin levels (Wangler et al. 2018). Since we observed reduced levels of intermediate chain length phospholipids in *Drosophila* in this study, we considered whether a related abnormality could underlie the sphingolipid reductions we observed in the human studies. Sphingomyelins are derived from phosphatidylcholine. However, the acyl chain in sphingomyelins are not derived from PC. Sphingomyelin receives the phosphocholine headgroup from PC and the carbon chains in sphingomyelin are derived from ceramide. Therefore, the precursor determining acyl chain length differs between sphingomyelin and phospholipids. Moreover, insects do not have prominent levels or biological roles for sphingomyelin (PC-ceramides). In insects such as *Drosophila*, PE-ceramides are of greater importance and abundance in insect membranes and, instead of an 18C-sphingoid base, contain a 14C sphingoid base(Rietveld et al. 1999). We therefore examined PE-ceramides in the fly brains. Examining across chain-lengths of PE ceramides we observed reduced quantities in *pex* mutants for PE Cer 16:0, PE Cer 18:0, PE Cer 20:0 and PE Cer 22:0 with increased PE Cer 24:0 and PE Cer 26:0 as reflected in the ratios for the mutant lines compared to control and rescue lines (**Figure 5A-B**). We observed a chain-length specific effect on PE ceramides in *Drosophila pex* mutant fly brain. PE Cer 22:0 is reduced in *pex2* mutants (**Figure 5C**) and in *pex16* mutants, although the hypomorphic *pex16^EY^* allele does not show this effect (**Figure 5D**). In contrast, longer chain length PE Cer 26:0 is increased in *pex2* mutants (**Figure 5E**) as well as *pex16* mutants (**Figure 5F**). In summary, we observe altered chain length of PE ceramides in the adult fly brain. This alteration is an imbalance toward longer chain lengths in fly brains.

**Figure 5.**
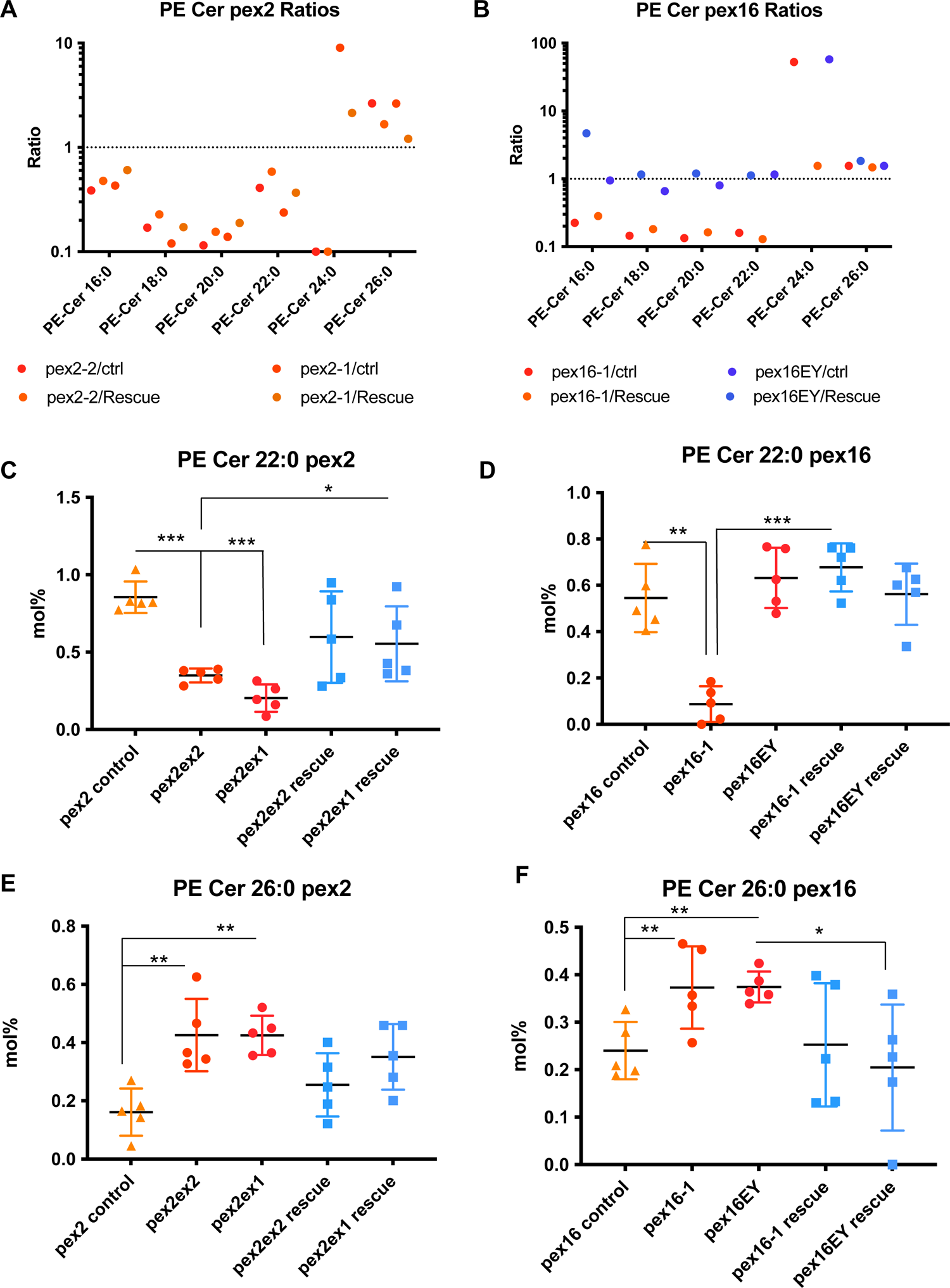
Chain-length specific PE ceramide abnormalities in peroxisomal mutant brain. **A.** Ratios of PE ceramides comparing *pex2^2^/Control* and *pex2^1^/Control* (red) and *pex2^2^/pex2^2^* rescue and *pex2^1^/pex2^1^* rescue (yellow). These ratios are notably lower for most PE ceramides, including 16:0-22:0, but the ratios are elevated for PE Cer 24:0 and PE Cer 26:0. **B.** Ratios of PE ceramides comparing *pex16^1^/Control* and *pex16^1^/Control* (red) and *pex16^EY^/pex16^EY^*rescue and *pex16^EY^/pex16^EY^* rescue (blue). These ratios show a pattern for *pex16^1^* with reduction in most PE ceramides, including 16:0-22:0, but the ratios are elevated for PE Cer 24:0 and PE Cer 26:0. **C.** Total levels in mol% of PE Cer 22:0 in *pex2* adult brains (genotypes shown in Figure 1) shows reduced levels in total PE Cer 22:0 in pex2 mutant brain compared to control (ratio *pex2^2/^control* 0.409, ratio *pex2^1/^control* 0.238, p<0.001, p<0.001 respectively), and shows reduced levels in total PE Cer 22:0 compared to pex2 rescue for *pex 2^1^* allele (ratio *pex2^1/^pex2^1^*rescue 0.368, p=0.028). **D.** Total levels in mol% of PE Cer 22:0 in *pex16* adult brains (genotypes shown in Figure 1) shows reduced levels in total PE Cer 22:0 in *pex16^1^*mutant brain compared to control (ratio *pex16^1/^control* 0.160, p=0.001), and *pex16^1^* mutant brain compared to rescue (ratio 0.129, p<0.001) but no significant differences were observed for *pex 16^EY^* allele. **E.** Total levels in mol% of PE Cer 26:0 in *pex2* adult brains (genotypes shown in Figure 1) shows increased levels in total PE Cer 26:0 in pex2 mutant brain compared to control (ratio *pex2^2/^control* 2.643, p=0.005, ratio *pex2^1/^control* 2.638, p<0.001, p=0.001 respectively), and increased on average compared to rescue although the differences were not significant. **F.** Total levels in mol% of PE Cer 26:0 in *pex16* adult brains (genotypes shown in Figure 1) shows increased levels in total PE Cer 26:0 in pex16 mutant brain compared to control (ratio *pex16^1/^control* 1.552, p=0.026, and increased fro *pex16^EY^* compared to control and to rescue (ratio *pex16^EY^*/control ratio 1.557, p=0.004, ratio *pex16^EY^*/rescue ratio 1.831, p=0.0439).

### Carbon chain length affects the levels of sphingomyelin in human subjects

Having observed a pattern of altered carbon chain lengths in *pex* mutant *Drosophila* we sought to confirm these observations in human samples. We examined the levels of PC, and sphingomyelin in human plasma samples. This included 16 individuals with PBD-ZSD due to *PEX1* genetic changes, the clinical details of these subjects has been previously reported (Wangler et al. 2018). We compared these samples to pediatric disease samples without peroxisomal disease from a distinct genetic cohort with no known peroxisomal link (Batzir et al. 2020), which we termed pediatric controls and we also tested unaffected adult controls. In plasma we observed increased relative levels of PC (**Figure 6A**). We also observed a decrease in the total levels of plasma ether-linked PC, i.e., alkyl or plasmenyl PC, here referred to as “ePC” (**Figure 6B**). Chain-length abnormalities were also observed, including decreased levels of PC 34:2 (**Figure 6C**), PC(36:2) (**Figure 6D**), PC 38:2 (**Figure 6E**), and PC 38:1 (**Figure 6F**) with an increase in the long chain PC 38:1 in patients with *PEX1* mutations compared to controls (**Figure 6B**). Increased levels of lysophosphatidylcholine were also observed in plasma (**Figure S10**). Indeed, a modest effect on PC molecular species according to chain length was observed in the human plasma samples with the PBD-ZSD patients exhibiting high levels of PC 40:3, PC 38:1, PC 38:2 and the low levels of PC 34:0 and PC 36:4 (**Figure S11**).

**Figure 6.**
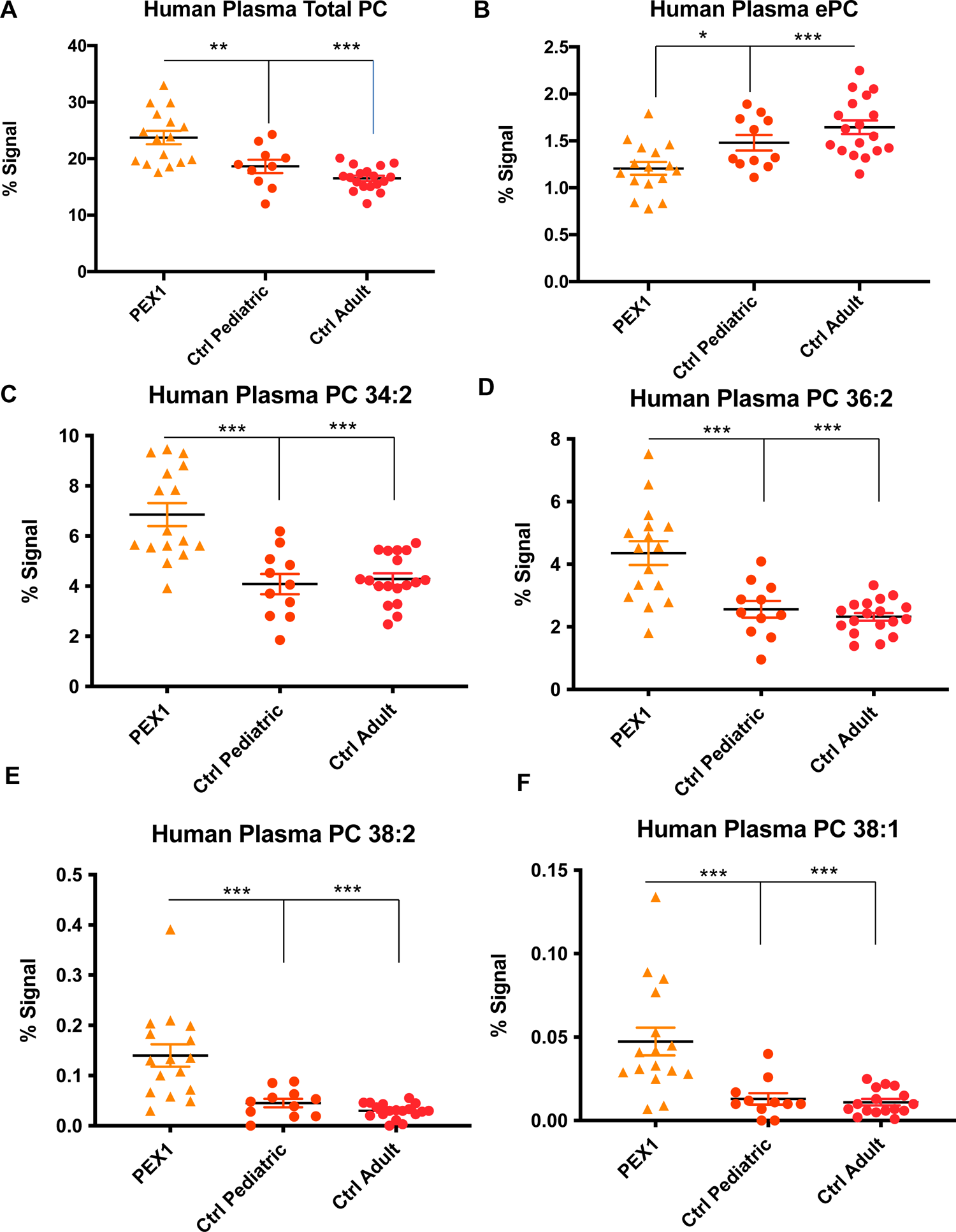
Phospholipid abnormalities in human plasma from patients with PBD-ZSD. **A.** Human plasma PC (as %) is increased in patients with PEX1 mutations compared to pediatric (ratio 1.26, p=0.005) and adult controls (ratio 1.437, p<0.001). **B.** Human plasma ether linked phosphatidylcholine (ePC) is reduced in patients with PEX1 mutations compared to pediatric (ratio 0.815, p=0.018) and adult controls (ratio 0.733, p<0.001). **C.** Human plasma PC 34:2 is increased in patients with PEX1 mutations compared to pediatric (ratio 1.678, p<0.001) and adult (ratio 1.600, p<0.001) controls. **D.** Human plasma PC 36:2 is increased in patients with PEX1 mutations compared to pediatric (ratio 1.702, p<0.001) and adult controls (ratio 1.878, p<0.001). **E.** Human plasma PC 38:2 is increased in patients with PEX1 mutations compared to pediatric (ratio 3.109, p<0.001) and adult controls (ratio 4.729, p<0.001). **F.** Human plasma PC 38:1 is increased in patients with PEX1 mutations compared to pediatric (ratio 3.679, p=0.001) and adult controls (ratio 3.784, p<0.001).

In sphingomyelin, we observed a reduction in SM 18:1 and SM 18:0 (**Figure 7A-B**) in patients with *PEX1* mutations. SM 22:0 was also reduced in the patients with PEX1 mutations (**Figure 7C**). However, for SM 24:0 the levels in plasma were the same for controls and the PBD-ZSD subjects (**Figure 7D**). Plasma DSM 22:0 was increased (**Figure 7E**). This pattern of abnormalities results in a dramatically increased ratio of C24/C22 sphingomyelins in patients with PBD-ZSD. In summary, there is a reduced level of certain subtypes of sphingomyelin in plasma with normal levels of very long chain sphingomyelins.

**Figure 7.**
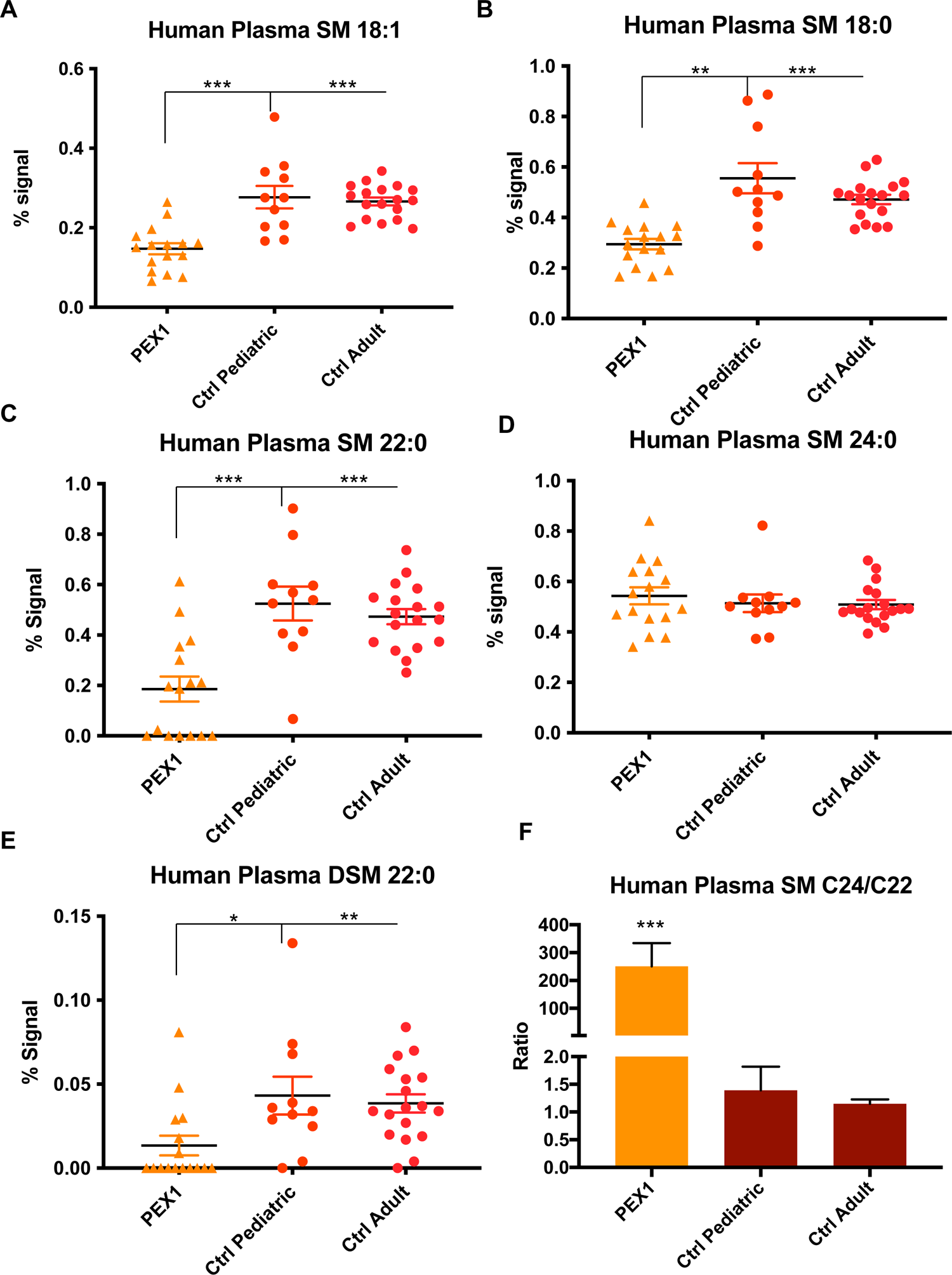
Sphingolipid abnormalities in human plasma from patients with PBD-ZSD. **A.** Human plasma SM 18:1 is decreased in patients with PEX1 mutations compared to pediatric (ratio 0.532, p<0.001) and adult controls (ratio 0.553, p<0.001). **B.** Human plasma SM 18:0 is decreased in patients with PEX1 mutations compared to pediatric (ratio 0.530, p=0.001) and adult controls (ratio 0.626, p<0.001). **C.** Human plasma SM 22:0 is decreased in patients with PEX1 mutations compared to pediatric (ratio 0.354, p=0.001) and adult controls (ratio 0.393, p<0.001). **D.** Human plasma SM 24:0 does not have differences for patients with PEX1 mutations compared to pediatric and adult controls. **E.** Human plasma dihydrosphingomyelin (DSM), 22:0 is reduced in patients with PEX1 mutations compared to pediatric (ratio 0.311, p=0.0327) and adult (ratio 0.349, p<0.001) controls **F.** The ratio of the human plasma SM C24/C22 shows a dramatic increase in the relative level of very long chain sphingomyelins in individuals with PEX1 mutations.

## Discussion

In this study we present a series of detailed lipidomic analyses on *Drosophila pex* mutants and human plasma samples from patients with PBD-ZSD. The principal findings of our study are that peroxisomal biogenesis mutations lead to reproducible alterations in phospholipids and sphingolipids in *Drosophila* larvae, brain and human plasma. These alterations have some consistent features in all these tissues including an overabundance of longer chain phospholipids like PC and PE and reduced relative amount of intermediate chain lengths. This unique *Drosophila* dataset is complementary with the findings related to human cells and tissues (Herzog et al. 2016, 2018). However, it also points to differences in the relative distribution of specific carbon chain-lengths, for example the overabundance of intermediate acyl chains in DAGs and their relative reduction in PEs and PCs. As DAGs are part of the Kennedy pathway for de novo synthesis of PEs and PCs, the imbalanced distribution suggests that regulation or effects such as substrate shuttling rather than simple chemical equilibrium are important factors to consider in PBD.

Defects in peroxisomal biogenesis have a well-documented overabundance of long and very long acyl chains as peroxisomes are required for the catabolism of very long chain fatty acids. In our dataset we observe increased long chain PCs and PEs but reduced long chain DAGs. The data are analyzed in mol% and therefore the overabundance of one analyte will lead to reduction in the percent of other analytes and so the increased long chain PCs and PEs are accompanied by reduced intermediate chain. However, for DAGs the inverse pattern, increased intermediate chains and reduced long chains are observed. PCs and PEs can be derived from diacyglycerol through the Kennedy pathway a process of *de novo* biosynthesis of phospholipids (Gibellini and Smith 2010). DAGs are thus converted to phospholipids via the Kennedy pathway. If these two pools of lipids (PEs and PCs versus DAGs) were in simple chemical equilibrium, the excess intermediate chain DAGs could be converted to intermediate chain PC and PEs and the long chain phospholipids could be converted to long chain DAGs and the imbalance we observe in the *pex* mutants would not exist. The fact that these imbalances are observed suggests that these lipid pools are not in equilibrium and this is an effect of substrate shuttling, a phenomenon in which certain substrates are preferentially incorporated into products in fatty acid pathways (Coleman 2019). The Kennedy pathway is not the only determinant of acyl chain length in phospholipids as specific acyl chains can be incorporated into PC and PE through exchange and remodeling via the Land’s cycle (Lands 1958; O’Donnell 2022). The Kennedy pathway for de novo biosynthesis through DAGs has preferential incorporation of the intermediate chain acyl chains into the resulting PEs and PCs due to preference of glycerol-3-phosphate acyltransferases (GPAT) (Lee and Ridgway 2020). However, the remodeling of the phospholipid through the Land’s cycle which exchanges acyl chains on the glycerol backbone has a preference for very long and long chains of some of the acyl transferases (O’Donnell 2022).

Applying our model, which takes into account substrate shuttling preferences, to our data leads us to propose a model for the phospholipid abnormalities in PBD (**Figure 8**). In our model the normal physiologic balance of acyl chains within PE, PC and DAGs and thus the mol% values in normal cells results from an overall balance in the acyl chain pool with some shuttling of in Kennedy pathway for intermediate chains and some shuttling of long chains in the Land’s cycle (**Figure 8A**). In PBD in contrast, the imbalance of acyl chains due to the peroxisomal fatty acid oxidation defect produces downstream phospholipid abnormalities that we observed (**Figure 8B**). The biochemical measurements at one point in time cannot provide information on the kinetics of enzymatic reactions in a pathway. It also cannot determine if an excess of long chain phosphatidylcholines is due to overproduction or lack of breakdown over time. However, it can reveal evidence for shuttling as 1) peroxisomes are required for very long chain fatty acid breakdown 2) increased levels of C24 and C26 derivatives of many biochemical lipids have been previously observed and 3) it seems likely that that is the proximal cause of the phospholipid alterations, and the changes observed in Intermediate chain lipids are the primary cause. Therefore, our model explains how the specific imbalances impact DAG’s (more influenced by the Kennedy pathway) and PE’s and PC’s (remodeled by Land’s cycle).

**Figure 8.**
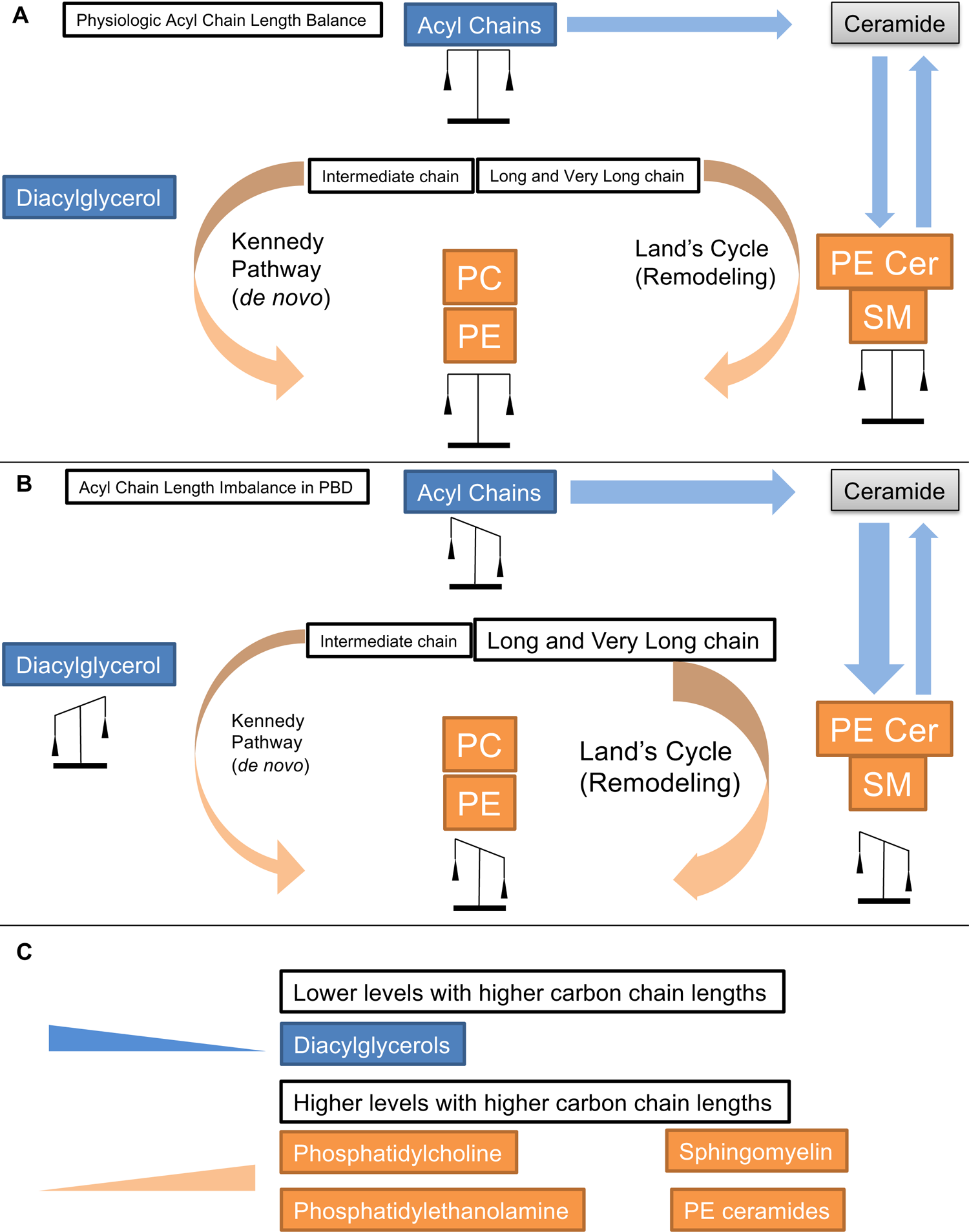
Biochemical model of phospholipid and sphingolipid abnormalities for peroxisomal biogenesis disorders. The Kennedy pathway mediates generation of phosphatidylcholine and phosphatidylethanolamine from diacylglycerol. Peroxisomal biogenesis defects lead to an excess of long and very long chain PC and PE, but an apparent reduction in intermediate chain. However the diacylglycerol levels appear to have an inverse relationship with reductions of very long and long chains and increased levels of intermediate chains. For sphingolipids (where the chain lengths are notably not in direct equilibrium with phospholipids) a similar excess of long and very long chains and reduced levels of intermediate chain. This loss of intermediate chains may explain the overall reduced levels of sphingomyelin observed in clinical samples.

Our model explains a number of observations we have previously been unable to account for. It explains why we had previously observed normal overall levels of PC and PE but unexplained increases in phosphocholine and phosphoethanolamine (Wangler et al. 2017) (**Figure S1**) as our model predicts that reduced flux through the Kennedy pathway would result. In addition, this data suggests an explanation for our report that sphingomyelin levels are reduced in plasma in PBD-ZSD (Wangler et al. 2018). We start with our data on PE-ceramides in *Drosophila* models in which we in fact observe a similar altered distribution of acyl chain length similar to what we see in PE and PC (**Figure 8**). In *Drosophila* and other insects PE ceramides are the predominant sphingolipid rather than Sphingomyelins (Vacaru et al. 2013). In these lipids the carbon chain lengths are derived from ceramides. We then extrapolate this to propose an explanation for the human sphingomyelin data (**Figure 7**). Similar to phospholipids the excess levels of very long chain lipids in peroxisomal mutants lead to overabundance of long chain sphingolipids such as sphingomyelin and PE ceramides and reduced overall levels of intermediate chain. This therefore explains the apparent reduced levels in the intermediate chains we previously reported (Wangler et al. 2018). Indeed, it was the *Drosophila* model allowed us to observe a dramatic overabundance of long chain PE ceramides and reduced levels of intermediate chains and to infer the cause of the clinical observations in humans. This is supported by our data, although we cannot rule out an overall deficiency of sphingomyelins with a disproportionate quantity of the remaining having long or very long chain acyl chains. Our analysis therefore points to significant alterations in the most common membrane lipids which we see in the brain. The defect in peroxisomal biogenesis that underlies PBD-ZSD likely has a dramatic impact on the composition of neuronal membranes, features that should be further explored for potential clinical consequences in PBD-ZSD.

This study provides a large dataset of phospholipid measurements in *Drosophila pex* mutants with some comparative studies in human plasma samples. Lipidomic analyses measuring phospholipids have been done in human cell lines and samples and there are areas of overlap within our study on human plasma samples that suggest the *Drosophila pex* mutants model many aspects of phospholipid biochemistry in PBD-ZSD. The *Drosophila* model allows us to systematically study the brain in adult flies and to provide observations from the brain *in vivo*.

## Acknowledgements

This work was supported by the National Institute for Neurological Disorders and Stroke 5R01NS107733 to MFW. The lipid analyses described in this work were performed at the Kansas Lipidomics Research Center Analytical Laboratory. Instrument acquisition and lipidomics method development were supported by the National Science Foundation (including support from the Major Research Instrumentation program; most recent award DBI-1726527), K-IDeA Networks of Biomedical Research Excellence (INBRE) of National Institute of Health (P20GM103418), USDA National Institute of Food and Agriculture (Hatch/Multi-State project 1013013), and Kansas State University.

## Supplementary Figure Legends

**Figure S1** Proposed phospholipid metabolomic alterations in *pex2* and *pex16 Drosophila* mutants from prior study In our previous metabolomic analysis of pex2 and pex16 mutants phospholipid abnormalities were identified which included a decrease in glycerol and glycerol-3-phosphate. Diacylglycerol is converted to phosphatidlycholine (with the addition of choline) or phosphatidylethanolamine (with the addition of ethanolamine) and the choline and ethanolamine precursors appear to be increased in the pex mutants. Also noted was an apparent excess of lysolipids and reduces levels of phospholipid breakdown products glycerophophorylcholine/glycerolethanolamine and glycerol 3-phosphate. Of note the levels of phosphatidylcholine and phosphatidylethanolamine themselves were normal, though the specific chain lengths were not analyzed in these previous studies.

**Figure S2 Robust chain-length specific reductions in intermediate chain phospholipids in peroxisomal mutant larvae** A. Total levels in mol% of PC 28:1 and PC 28:0 Phosphatidylcholine (PC) in *pex2* and *pex16* larvae shows dramatic and significant decreases in intermediate chain phospholipids. For PC 28:1, *pex2* mutant larvae compared to *pex2* rescue (ratio *pex2/pex2* rescue: ratio 0.1889, p=0.0006), and a dramatic decrease in *pex16* mutant larvae compared to *pex16* rescue (*pex16/pex16* rescue: ratio 0.211, p=0.003). For PC 28:0, *pex2* mutant larvae compared to *pex2* rescue (ratio *pex2/pex2* rescue: ratio 0.427,p=0.0002), and a non-significant decrease in *pex16* mutant larvae compared to *pex16* rescue (*pex16/pex16* rescue: ratio 0.502, p=0.0555). B. Total levels in mol% of PE 28:1 and PE 28:0 Phosphatidylethanolamine (PE) in *pex2* and *pex16* larvae shows dramatic and significant decreases in intermediate chain phospholipids. For PE 28:1, *pex2* mutant larvae compared to *pex2* rescue (ratio *pex2/pex2* rescue: ratio 0.2369, p=0.0003), and a dramatic decrease in *pex16* mutant larvae compared to *pex16* rescue (*pex16/pex16* rescue: ratio 0.233, p=7.97E-05). For PE 28:0, *pex2* mutant larvae compared to *pex2* rescue (ratio *pex2/pex2* rescue: ratio 0.594, p=0.0027), and a non-significant decrease in *pex16* mutant larvae compared to *pex16* rescue (*pex16/pex16* rescue: ratio 0.624, p=0.0022). C. Total levels in mol% of PC 30:2, PC 30:0, PC 31:2, PC31:1, and PC 31:0 Phosphatidylcholine (PC) in *pex2* and *pex16* larvae showing dramatic and significant decreases in intermediate chain phospholipids. For PC 30:2, *pex2* mutant larvae compared to *pex2* rescue (ratio *pex2/pex2* rescue: ratio 0.1545, p=0.0002), and a dramatic decrease in *pex16* mutant larvae compared to *pex16* rescue (*pex16/pex16* rescue: ratio 0.1775, p=0.0018). For PC 30:0, *pex2* mutant larvae compared to *pex2* rescue (ratio *pex2/pex2* rescue: ratio 0.5748, p=0.0303), and a dramatic decrease in *pex16* mutant larvae compared to *pex16* rescue (*pex16/pex16* rescue: ratio 0.6542, p=0.3048). For PC 31:2, *pex2* mutant larvae compared to *pex2* rescue (ratio *pex2/pex2* rescue: ratio 0.1857, p=0.0020), and a dramatic decrease in *pex16* mutant larvae compared to *pex16* rescue (*pex16/pex16* rescue: ratio 0.2664, p=0.0427). For PC 31:1, *pex2* mutant larvae compared to *pex2* rescue (ratio *pex2/pex2* rescue: ratio 0.3812, p=0.0003), and a dramatic decrease in *pex16* mutant larvae compared to *pex16* rescue (*pex16/pex16* rescue: ratio 0.5449, p=0.0150). For PC 31:0, *pex2* mutant larvae compared to *pex2* rescue (ratio *pex2/pex2* rescue: ratio 0.5149, p=0.0002), and a non-significant decrease in *pex16* mutant larvae compared to *pex16* rescue (*pex16/pex16* rescue: ratio 0.6767, p=0.2111).

**Figure S3** Robust chain-length specific elevations in long chain phospholipids in peroxisomal mutant larvae Total levels in mol% of PC 37:2: and PC 38:2 Phosphatidylcholine (PC) in *pex2* and *pex16* larvae shows dramatic and significant increases in long chain phospholipids. For PC 37:2, *pex2* mutant larvae compared to *pex2* rescue (ratio *pex2/pex2* rescue: ratio 2.473, p=0.0183), and a dramatic increase in *pex16* mutant larvae compared to *pex16* rescue (*pex16/pex16* rescue: ratio 3.446, p=0.002). For PC 38:2, *pex2* mutant larvae compared to *pex2* rescue (ratio *pex2/pex2* rescue: ratio 2.303, p=0.0426), and a non-significant decrease in *pex16* mutant larvae compared to *pex16* rescue (*pex16/pex16* rescue: ratio 3.243, p=0.0193).

**Figure S4** Increasing levels of PCs based on chain length across different classes of lipid side chains organized by unsaturations The same ratios depicted in Figure 3A is reorganized based on the total number of unsaturations, organized from (N:0) up to (N:4) showing that across each class the ratios or pex2/pex2 rescue and pex16/pex16 rescue increases as chain-length increases. Clear trends are not observed for the number of unsaturations themselves.

**Figure S5** Robust chain-length specific reductions in intermediate chain phospholipids in peroxisomal mutant brain **A.** Levels in mol% of PC 30:1 in *pex2* and *pex16* mutant brains shows significant reduction in *pex* mutant brains. For PC 30:1 in *pex2*^2^ brain and *pex2^1^* brain, levels are reduced compared to controls (ratio 0.733, 0.810 respectively, p=0.014, p=0.049 respectively) and *pex2^2^*brain compared to rescue (ratio 0.856, p=0.028), with no difference for *pex2^1^* compared to rescue (ratio 0.987, p=0.764). For PC 30:1 in *pex16^1^* and *pex16^EY^* brain, levels are reduced compared to controls (ratio 0.675, 0.826 respectively, p<0.001, p=0.028 respectively). And *pex16^1^* brain compared to rescue (ratio 0.832, p=0.014), with no difference for *pex16^EY^* compared to rescue (ratio 1.046, p=0.679). **B.** Levels in mol% of PC 30:2 in *pex2* and *pex16* mutant brains shows significant reduction in *pex* mutant brains. For PC 30:2 in *pex2*^2^ brain and *pex2^1^* brain, levels are reduced compared to controls (ratio 0.526, 0.575 respectively, p=0.004, p=0.004 respectively) and *pex2^2^*brain compared to rescue (ratio 0.655, p<0.001) as well as *pex2^1^* compared to rescue (ratio 1.021, p=0.837). For PC 30:2 in *pex16^1^* and *pex16^EY^* brain, levels are reduced compared to controls (ratio 0.387, 0.624 respectively, p<0.001, p=0.008 respectively). And *pex16^1^* brain compared to rescue (ratio 0.525, p=0.002), with no difference for *pex16^EY^* compared to rescue (ratio 1.021, p=0.837).

**Figure S6** Robust chain-length specific elevations in some intermediate chain and long chain phospholipids in peroxisomal mutant brain **A.** Levels in mol% of PC 30:0 in *pex2* and *pex16* mutant brains show significant increases in *pex* mutant brains. For PC 30:0 in *pex2*^2^ brain and *pex2^1^* brain, levels are increased compared to controls (ratio 1.131, 1.238 respectively, p=0.038, p=0.001 respectively) and *pex2^2^*brain compared to rescue (ratio 1.383, p<0.001), and *pex2^1^*compared to rescue (ratio 1.483, p<0.001). For PC 30:0 in *pex16^1^* and *pex16^EY^* brain, levels are increased compared to controls (ratio 1.368, 1.107 respectively, p<0.001, p=0.001 respectively). And *pex16^1^* brain compared to rescue (ratio 1.186, p=0.002), and increased *pex16^EY^* compared to rescue (ratio 1.222, p=0.008). **B.** Levels in mol% of PC 35:1 in *pex2* and *pex16* mutant brains shows significant increases in *pex* mutant brains. For PC 35:1 in *pex2*^2^ brain and *pex2^1^* brain, levels are increased compared to controls (ratio 2.529, 1.653 respectively, p=0.003, p=0.004 respectively) and *pex2^2^*brain compared to rescue (ratio 1.985, p=0.005) as well as *pex2^1^* compared to rescue (ratio 2.059, p=0.006). For PC 35:1 in *pex16^1^* and *pex16^EY^* brain, levels are increased compared to controls (ratio 2.129, 2.310 respectively, p<0.001, p<0.001 respectively). And *pex16^1^* brain compared to rescue (ratio 1.629, p=0.019), and for *pex16^EY^* compared to rescue (ratio 1.690, p<0.001). **C.** Levels in mol% of PC 34:1 in *pex2* and *pex16* mutant brains shows significant increases in *pex* mutant brains. For PC 34:1 in *pex2*^2^ brain and *pex2^1^* brain, levels are increased compared to controls (ratio 1.387, 1.407 respectively, p<0.001, p<0.001 respectively) and *pex2^2^* brain compared to rescue (ratio 1.289, p<0.001) as well as *pex2^1^* compared to rescue (ratio 1.365, p<0.001). For PC 34:1 in *pex16^1^* levels are increased compared to controls (ratio 1.137, respectively, p<0.001, p<0.001 respectively) and rescue (ratio 1.321, p=0.002). However for *pex16^EY^*brain the levels are mildly reduced compared to control (ratio 0.898, p=0.006), and *pex16^EY^* compared to rescue (ratio 0.885, p=0.017). **D.** Levels in mol% of PC 34:2 in *pex2* and *pex16* mutant brains shows significant increases in *pex* mutant brains. For PC 34:2 in *pex2*^2^ brain and *pex2^1^* brain, levels are increased compared to controls (ratio 1.217, 1.226 respectively, p=0.002, p<0.001 respectively) and *pex2^2^* brain compared to rescue (ratio 1.184, p=0.005) as well as *pex2^1^* compared to rescue (ratio 1.188, p=0.001). For PC 34:2 in *pex16^1^* levels are increased compared to controls (ratio 1.092, p=0.031) and compared to rescue (ratio 1.218, p=0.001). However, *pex16^EY^*brain has reduced levels compared to controls (ratio 0.895, p=0.012). and no difference from rescue. **E.** Levels in mol% of PC 37:2 in *pex2* and *pex16* mutant brains shows significant increases in *pex* mutant brains. For PC 37:2 in *pex2*^2^ brain and *pex2^1^*brain, levels are increased compared to controls (ratio 2.471, 1.939 respectively, p=0.001, p=0.004 respectively) and *pex2^2^* brain compared to rescue (ratio 2.722, p=0.001) as well as *pex2^1^* compared to rescue (ratio 1.806, p=0.008). For PC 37:2 in *pex16^1^* and *pex16^EY^* brain levels are increased compared to controls (ratio 1.860, 1.871 respectively, p=0.005, p=0.003 respectively) and *pex16^1^* compared to rescue (ratio 1.755, p=0.009) and for *pex16^EY^* compared to rescue (ratio 2.001, p=0.009). **F.** Levels in mol% of PC 37:1 in *pex2* and *pex16* mutant brains shows significant increases in *pex* mutant brains. For PC 37:1 in *pex2*^2^ brain and *pex2^1^*brain, levels are increased compared to controls (ratio 3.770, 2.504 respectively, p=0.001, p=0.010 respectively) and *pex2^2^* brain compared to rescue (ratio 2.621, p=0.003) as well as *pex2^1^* compared to rescue (ratio 1.628, p=0.022). For PC 37:1 in *pex16^1^* and *pex16^EY^* brain levels are increased compared to controls (ratio 2.703, 2.442 respectively, p<0.001, p<0.001 respectively) and *pex16^1^* compared to rescue (ratio 1.519, p=0.003) and non-significant increase for *pex16^EY^* compared to rescue (ratio 1.398, p=0.065).

**Figure S7** Robust chain-length specific reductions in some intermediate chain and long chain PE phospholipids in peroxisomal mutant brain **A.** Levels in mol% of PE 33:3 in *pex2* and *pex16* mutant brains show significant decreases in *pex* mutant brains. For PE 33:3 in *pex2*^2^ brain and *pex2^1^* brain, levels are dramatically reduced compared to controls (ratio 0.225, 0.200 respectively, p<0.001, p<0.001 respectively) and *pex2^2^* brain compared to rescue (ratio 0.244, p<0.001), and *pex2^1^* compared to rescue (ratio 0.220, p<0.001). For PE 33:3 in *pex16^1^* and *pex16^EY^* brain, levels are decreased compared to controls (ratio 0.214, 0.718 respectively, p<0.001, p<0.001 respectively). And *pex16^1^* brain compared to rescue (ratio 0.249, p=0.002), and for *pex16^EY^* compared to rescue (ratio 0.756, p<0.001). **B.** Levels in mol% of PE 33:2 in *pex2* and *pex16* mutant brains show significant decreases in *pex* mutant brains. For PE 33:2 in *pex2*^2^ brain and *pex2^1^* brain, levels are dramatically reduced compared to controls (ratio 0.403, 0.469 respectively, p<0.001, p<0.001 respectively) and *pex2^2^* brain compared to rescue (ratio 0.430, p<0.001), and *pex2^1^* compared to rescue (ratio 0.496, p<0.001). For PE 33:2 in *pex16^1^* and *pex16^EY^* brain, levels are decreased compared to controls (ratio 0.475, 0.688 respectively, p<0.001, p<0.001 respectively). And *pex16^1^* brain compared to rescue (ratio 0.494, p=0.006), and for *pex16^EY^* compared to rescue (ratio 0.658, p=0.001). **C.** Levels in mol% of PE 33:1 in *pex2* and *pex16* mutant brains show significant decreases in *pex* mutant brains. For PE 33:3 in *pex2*^2^ brain and *pex2^1^* brain, levels are dramatically reduced compared to controls (ratio 0.549, 0.578 respectively, p<0.001, p<0.001 respectively) and *pex2^2^* brain compared to rescue (ratio 0.528, p<0.001), and *pex2^1^*compared to rescue (ratio 0.510, p<0.001). For PE 33:3 in *pex16^1^* levels are decreased compared to controls (ratio 0.789, p<0.001). And *pex16^1^* brain compared to rescue (ratio 0.758, p=0.019). For the *pex16^EY^* the difference was not significant from control, but was reduced compared to rescue (ratio 0.868, p=0.006). **D.** Levels in mol% of PE 35:4 in *pex2* and *pex16* mutant brains show significant decreases in *pex* mutant brains. For PE 35:4 in *pex2*^2^ brain and *pex2^1^* brain, levels are dramatically reduced compared to controls (ratio 0.400, 0.467 respectively, p<0.001, p<0.001 respectively) and *pex2^2^* brain compared to rescue (ratio 0.380, p<0.001), and *pex2^1^*compared to rescue (ratio 0.528, p<0.001). For PE 35:4 in *pex16^1^* levels are decreased compared to controls (ratio 0.443, p=0.002). And *pex16^1^* brain compared to rescue (ratio 0.433, p=0.047). For the *pex16^EY^* the difference was not significant from control, but was reduced compared to rescue (ratio 0.749, p<0.001). **E.** Levels in mol% of PE 35:3 in *pex2* and *pex16* mutant brains show significant decreases in *pex* mutant brains. For PE 35:3 in *pex2*^2^ brain and *pex2^1^* brain, levels are dramatically reduced compared to controls (ratio 0.131, 0.119 respectively, p<0.001, p<0.001 respectively) and *pex2^2^* brain compared to rescue (ratio 0.110, p<0.001), and *pex2^1^* compared to rescue (ratio 0.101, p<0.001). For PE 35:3 in *pex16^1^* and *pex16^EY^* levels are decreased compared to controls (ratio 0.141, 0.644, p<0.001, P<0.001) and for *pex16^1^* brain compared to rescue (ratio 0.181, p=0.001) and for *pex16^EY^* compared to rescue (ratio 0.677, p<0.001). **F.** Levels in mol% of PE 35:2 in *pex2* and *pex16* mutant brains show significant decreases in *pex* mutant brains. For PE 35:2 in *pex2*^2^ brain and *pex2^1^* brain, levels are dramatically reduced compared to controls (ratio 0.118, 0.124, respectively, p<0.001, p<0.001 respectively) and *pex2^2^* brain compared to rescue (ratio 0.104, p<0.001), and *pex2^1^* compared to rescue (ratio 0.115, p<0.001). For PE 35:2 in *pex16^1^* and *pex16^EY^* levels are decreased compared to controls (ratio 0.186, 0.540, p<0.001, P<0.001) and for *pex16^1^* brain compared to rescue (ratio 0.227, p=0.006) and for *pex16^EY^* compared to rescue (ratio 0.527, p<0.001).

**Figure S8** Robust chain-length specific elevations in some intermediate chain and long chain PE phospholipids in peroxisomal mutant brain **A.** Levels in mol% of PE 33:0 in *pex2* and *pex16* mutant brains show significant increases in *pex* mutant brains. For PE 33:0 in *pex2*^2^ brain and *pex2^1^* brain, levels are increased compared to controls (ratio 2.227, 2.153 respectively, p<0.001, p=0.009 respectively) and *pex2^2^* brain compared to rescue (ratio 2.032, p<0.001), and *pex2^1^* compared to rescue (ratio 1.644, p=0.044). For PE 33:0 in *pex16^1^* and *pex16^EY^* brain, levels are increased compared to controls (ratio 1.834, 1.900 respectively, p=0.005, p=0.001 respectively) and for *pex16^EY^* compared to rescue (ratio 1.518, p=0.003). **B.** Levels in mol% of PE 34:4 in *pex2* and *pex16* mutant brains show significant decreases in *pex* mutant brains. For PE 34:4 in *pex2*^2^ brain and *pex2^1^* brain, levels are increased compared to controls (ratio 1.285, 1.202 respectively, p=0.001, p=0.005 respectively) and *pex2^2^* brain compared to rescue (ratio 1.151, p=0.002). For PE 34:4 in *pex16^1^* and *pex16^EY^* brain, levels are increased compared to controls (ratio 1.366, 1.136 respectively, p<0.001, p=0.048 respectively) but no differences from rescue. **C.** Levels in mol% of PE 34:3 in *pex2* and *pex16* mutant brains show significant increases in *pex* mutant brains. For PE 34:3 in *pex2*^2^ brain and *pex2^1^* brain, levels are increased compared to controls (ratio 1.327, 1.343 respectively, p<0.001, p<0.001 respectively) and *pex2^2^* brain compared to rescue (ratio 1.170, p<0.001), and *pex2^1^* compared to rescue (ratio 1.094, p=0.002). For PE 34:3 in *pex16^1^* and *pex16^EY^* brain, levels are increased compared to controls (ratio 1.388, 1.157 respectively, p<0.001, p<0.001 respectively) and for *pex16^1^* compared to rescue (ratio 1.141, p=0.003). **D.** Levels in mol% of PE 34:1 in *pex2* and *pex16* mutant brains show significant increases in *pex* mutant brains. For PE 34:1 in *pex2*^2^ brain and *pex2^1^* brain, levels are increased compared to controls (ratio 1.371, 1.328 respectively, p<0.001, p<0.001 respectively) and *pex2^2^* brain compared to rescue (ratio 1.219, p<0.001), and *pex2^1^* compared to rescue (ratio 1.141, p<0.001). For PE 34:1 in *pex16^1^* and *pex16^EY^* brain, levels are increased compared to controls (ratio 1.249, 1.140 respectively, p<0.001, p<0.001 respectively) and for *pex16^EY^* compared to rescue (ratio 1.076, p<0.001). **E.** Levels in mol% of PE 36:3 in *pex2* and *pex16* mutant brains show significant increases in *pex* mutant brains. For PE 36:3 in *pex2*^2^ brain and *pex2^1^* brain, levels are increased compared to controls (ratio 1.180, 1.240 respectively, p<0.001, p<0.001 respectively) and *pex2^2^* brain compared to rescue (ratio 1.056, p=0.025), and *pex2^1^* compared to rescue (ratio 1.068, p=0.009). For PE 36:3 in *pex16^1^* and *pex16^EY^*brain, levels are increased compared to controls (ratio 1.112, 1.157 respectively, p<0.001, p<0.001 respectively) and for *pex16^EY^* compared to rescue (ratio 1.092, p=0.002). **F.** Levels in mol% of PE 38:3 in *pex2* and *pex16* mutant brains show significant increases in *pex* mutant brains. For PE 38:3 in *pex2*^2^ brain and *pex2^1^* brain, levels are increased compared to controls (ratio 1.795, 1.717 respectively, p<0.001, p<0.001 respectively) and *pex2^2^* brain compared to rescue (ratio 1.661, p<0.001), and *pex2^1^* compared to rescue (ratio 1.540, p<0.001). For PE 38:3 in *pex16^1^* and *pex16^EY^* brain, levels are increased compared to controls (ratio 1.393, 1.382 respectively, p<0.001, p=0.001 respectively) and for *pex16^1^* compared to rescue (ratio 1.201, p=0.005) *pex16^EY^* compared to rescue (ratio 1.157, p=0.029).

**Figure S9** Complex relationship between phospholipid ratios spanning intermediate and long chain lengths in adult fly brain A. Ratios of PCs comparing *pex2^1^/ control* (red) and *pex2^1^/pex2^1^* rescue (yellow). These ratios may show some mild increase as the chain length increase but there is variability. B. Ratios of PCs comparing *pex16^1^/ control* (red) and *pex16^1^/pex16^1^* rescue (yellow). These ratios may show some mild increase as the chain length increase but there is variability. C. Ratios of PCs comparing *pex16^EY^/ control* (blue) and *pex16^EY^/pex16^1^* rescue (blue). These ratios show no clear pattern depending on chain length.

**Figure S10** Lyso-Phospholipid abnormalities in human plasma from patients with PBD-ZSD **A.** Human Plasma LPC(16:0) is decreased in patients with PEX1 mutations compared to pediatric (ratio 0.523, p<0.001) and adult controls (ratio 0.656, p<0.001). **B.** Human Plasma LPC(18:0) is decreased in patients with PEX1 mutations compared to pediatric (ratio 0.467, p<0.001) and adult controls (ratio 0.620, p=0.001).

**Figure S11** Ratio across chain-length for phospholipids in human plasma samples for patients with PEX1 mutations **A.** Ratios of PCs comparing *PEX1 samples/Pediatric controls* (red) and *PEX1 samples/Adult controls* (red) (yellow). **B.** Ratios shown in panel A reorganized according to the number of unsaturations (N).

