## Supplementary figures and images for "Drosophila Models Uncover Substrate Channeling Effects on Phospholipids and Sphingolipids in Peroxisomal Biogenesis Disorders"

### Figure S1

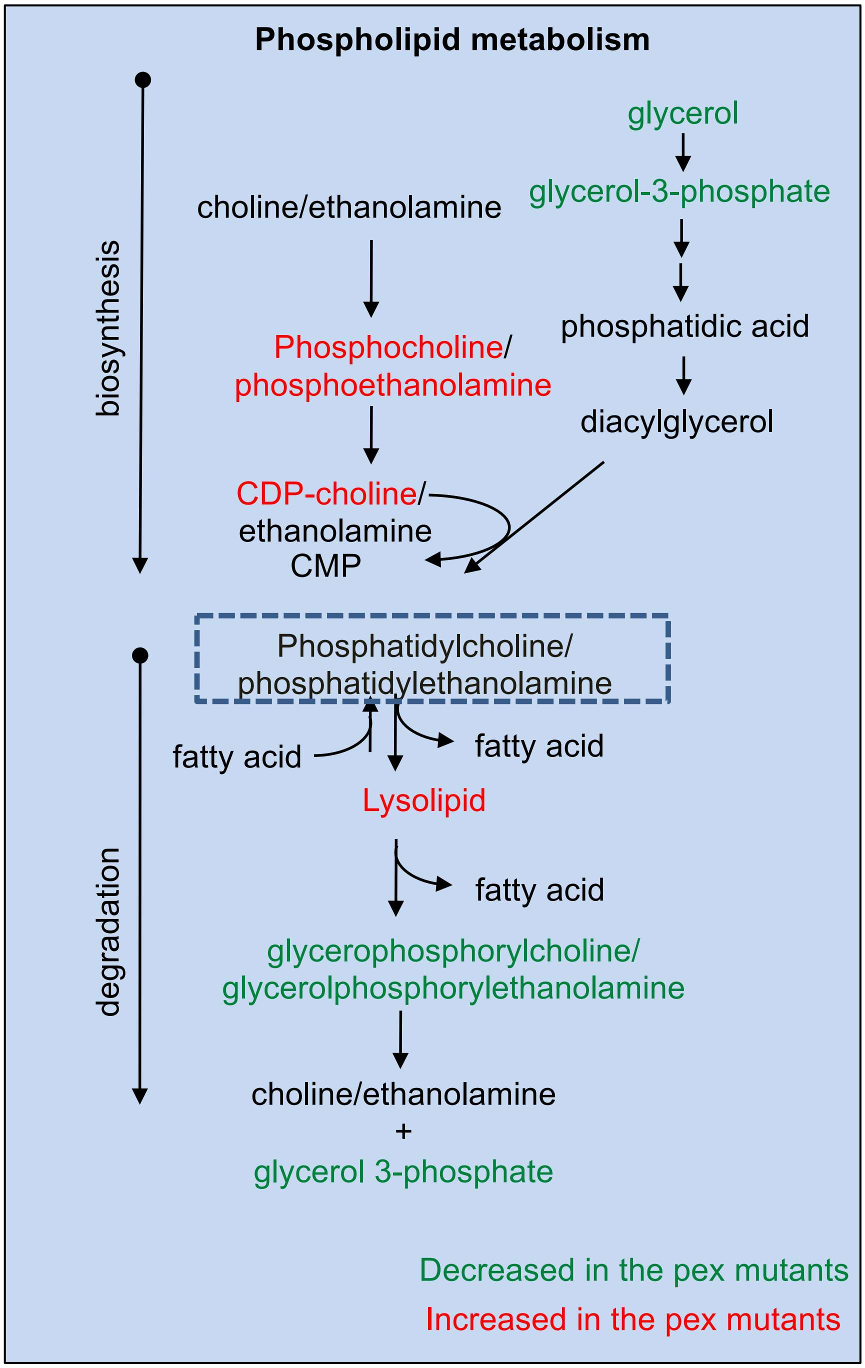

### Figure S2

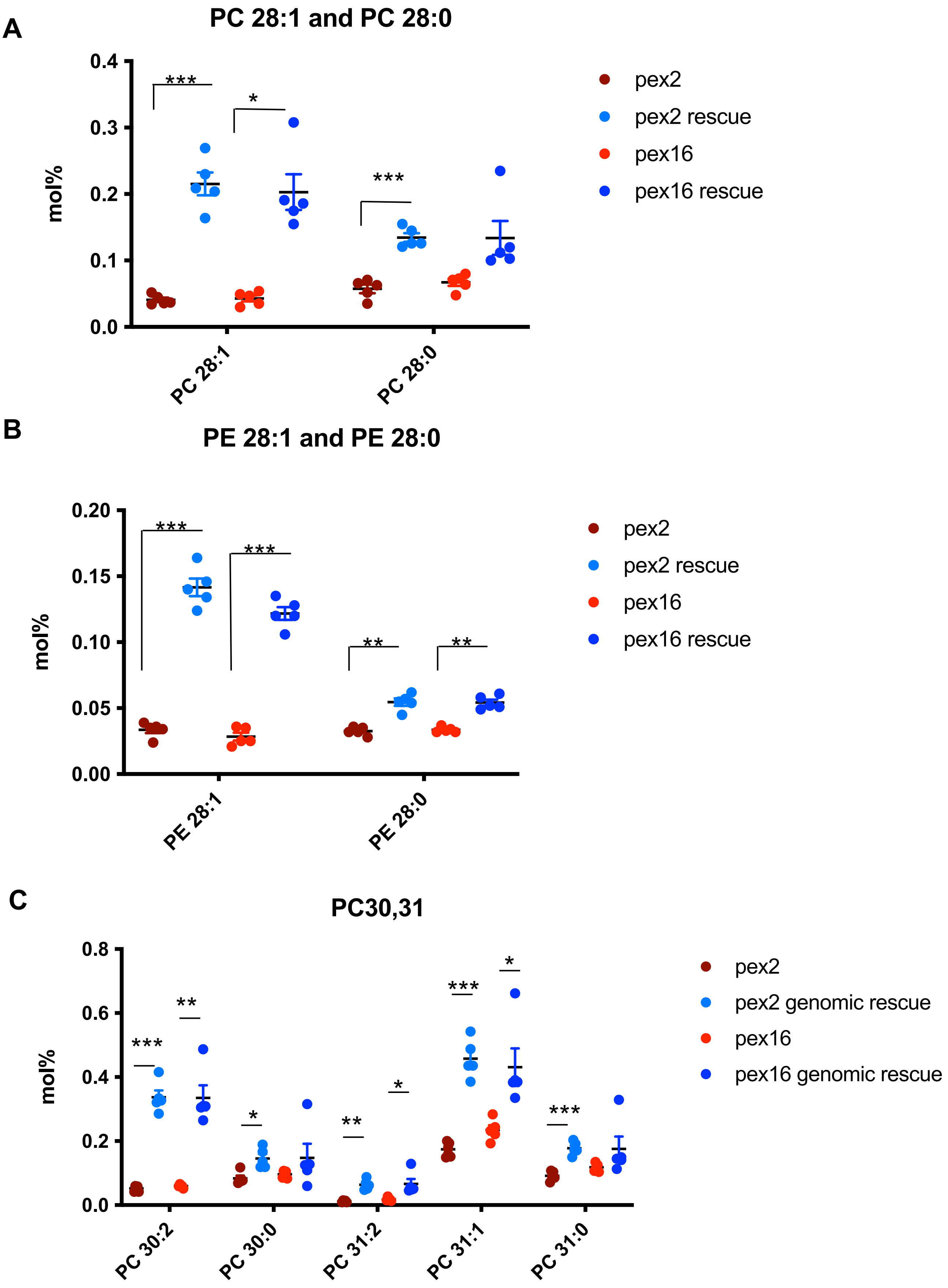

### Figure S3

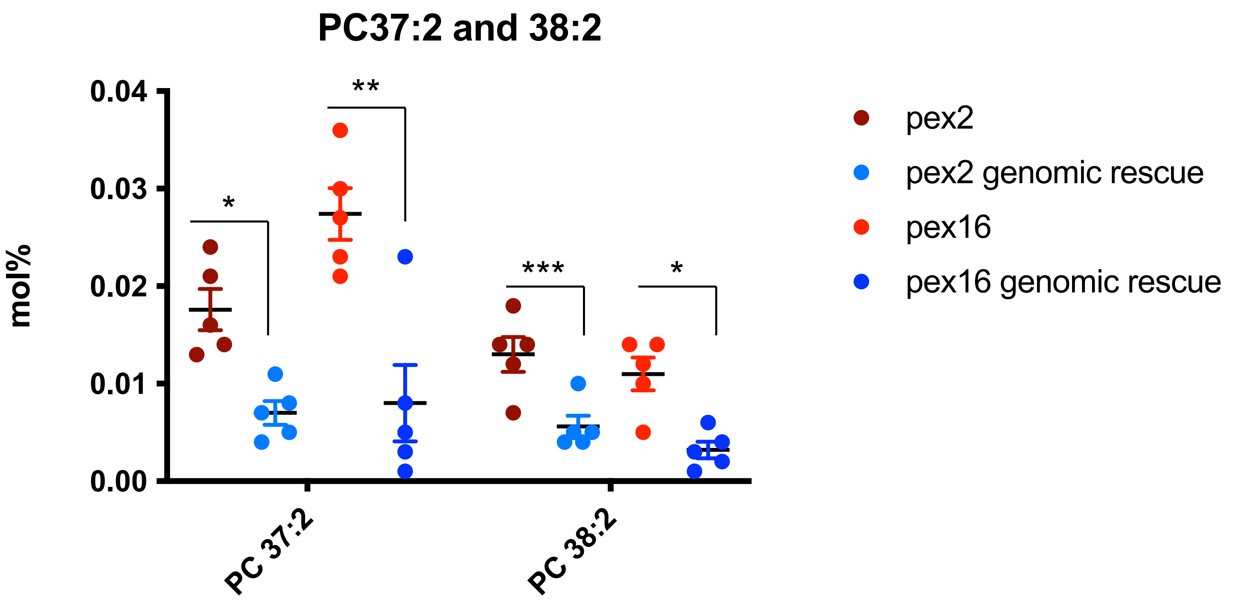

### Figure S4

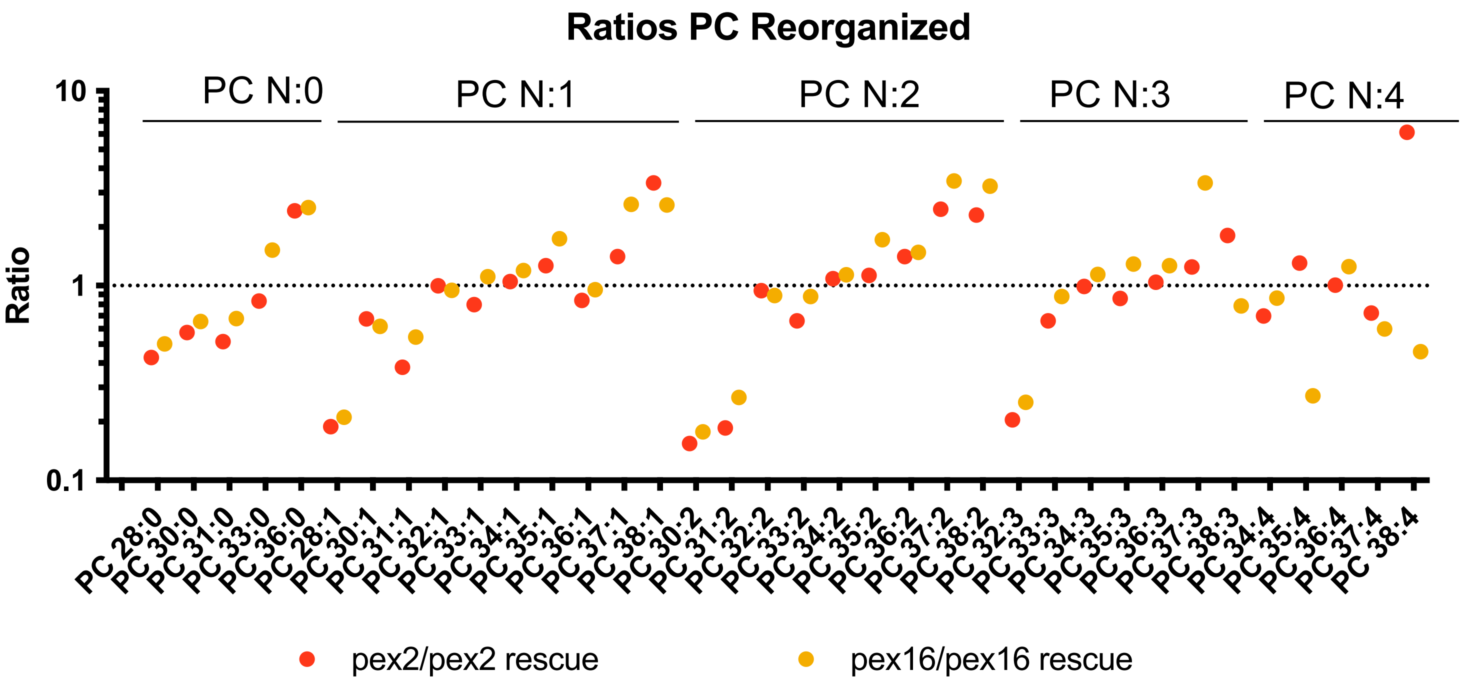

### Figure S5

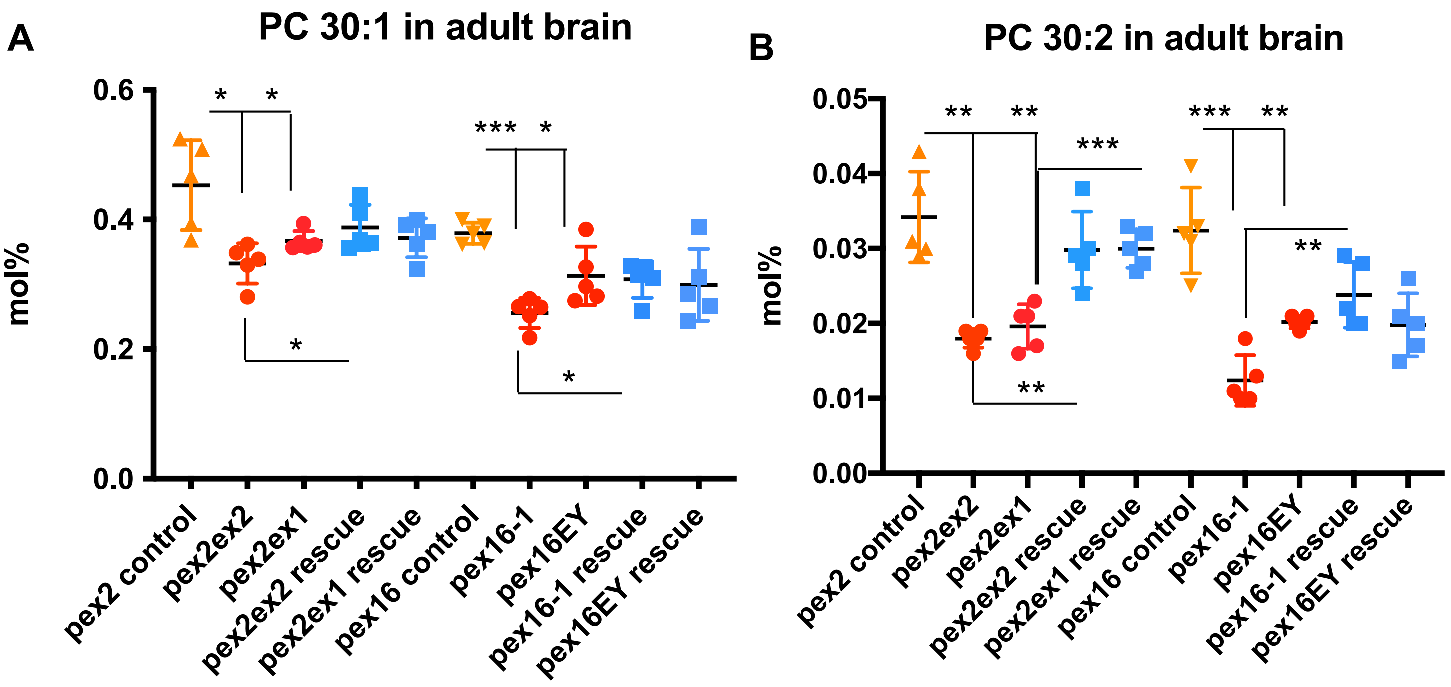

### Figure S6

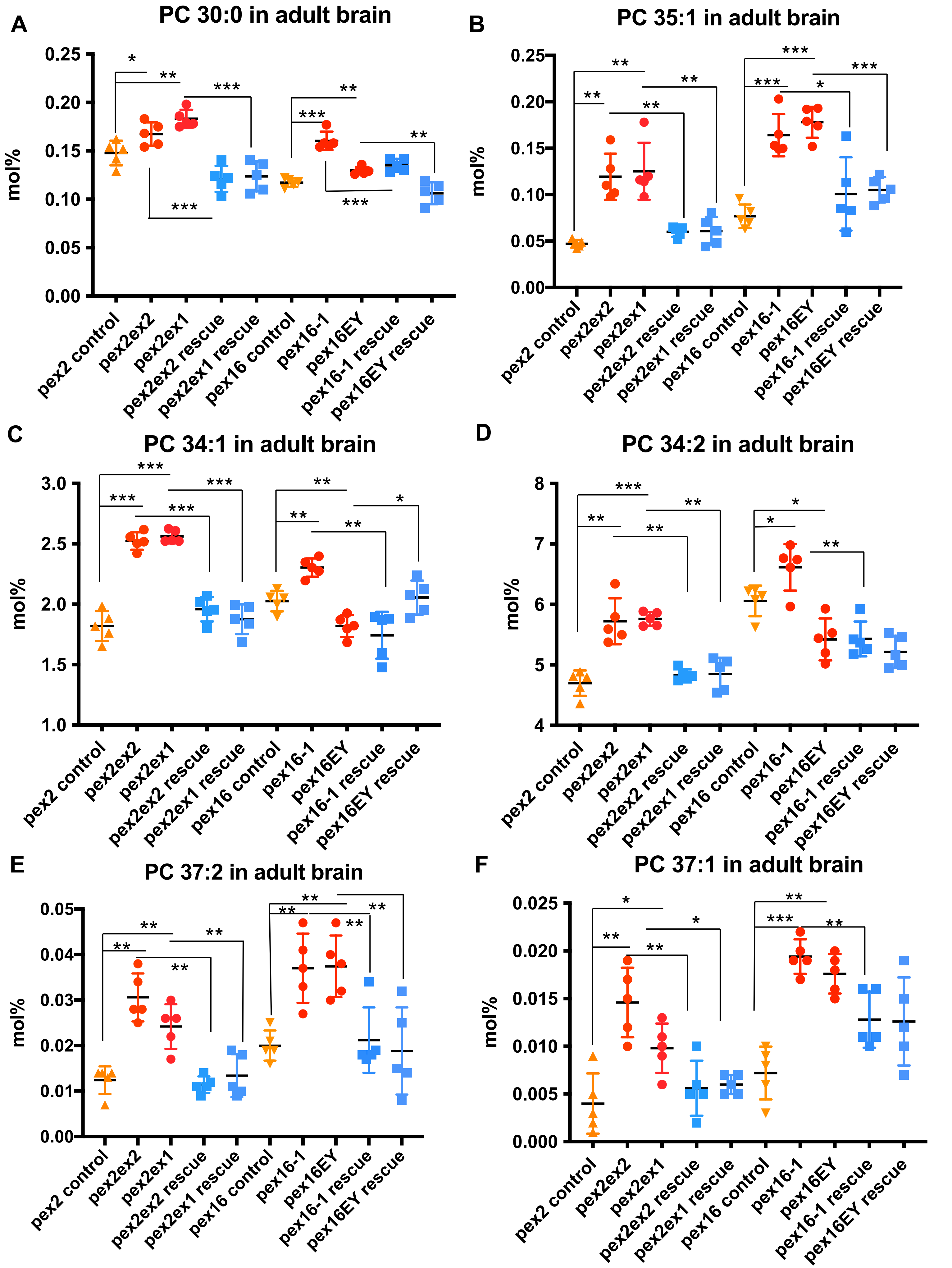

### Figure S7

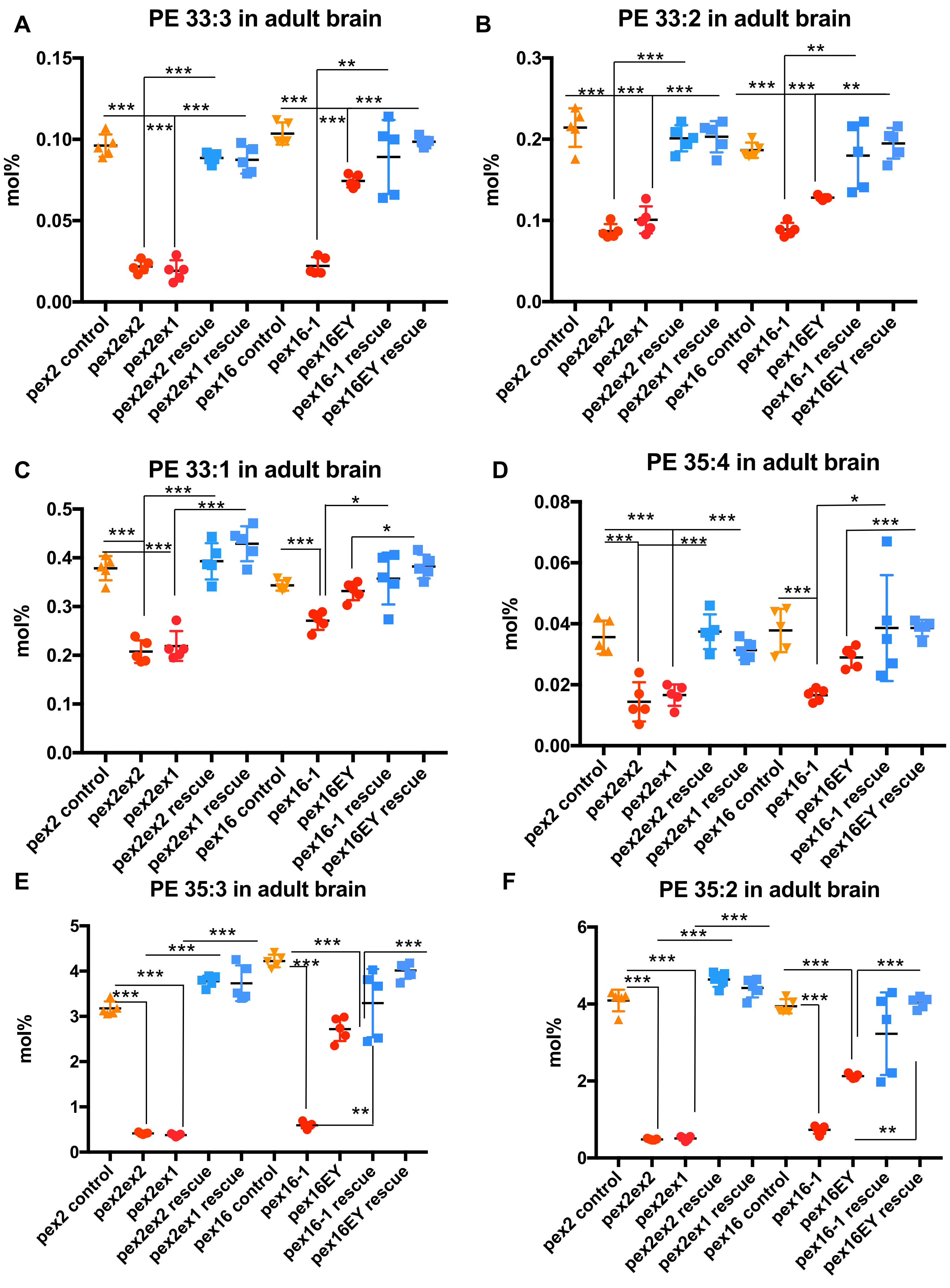

### Figure S8

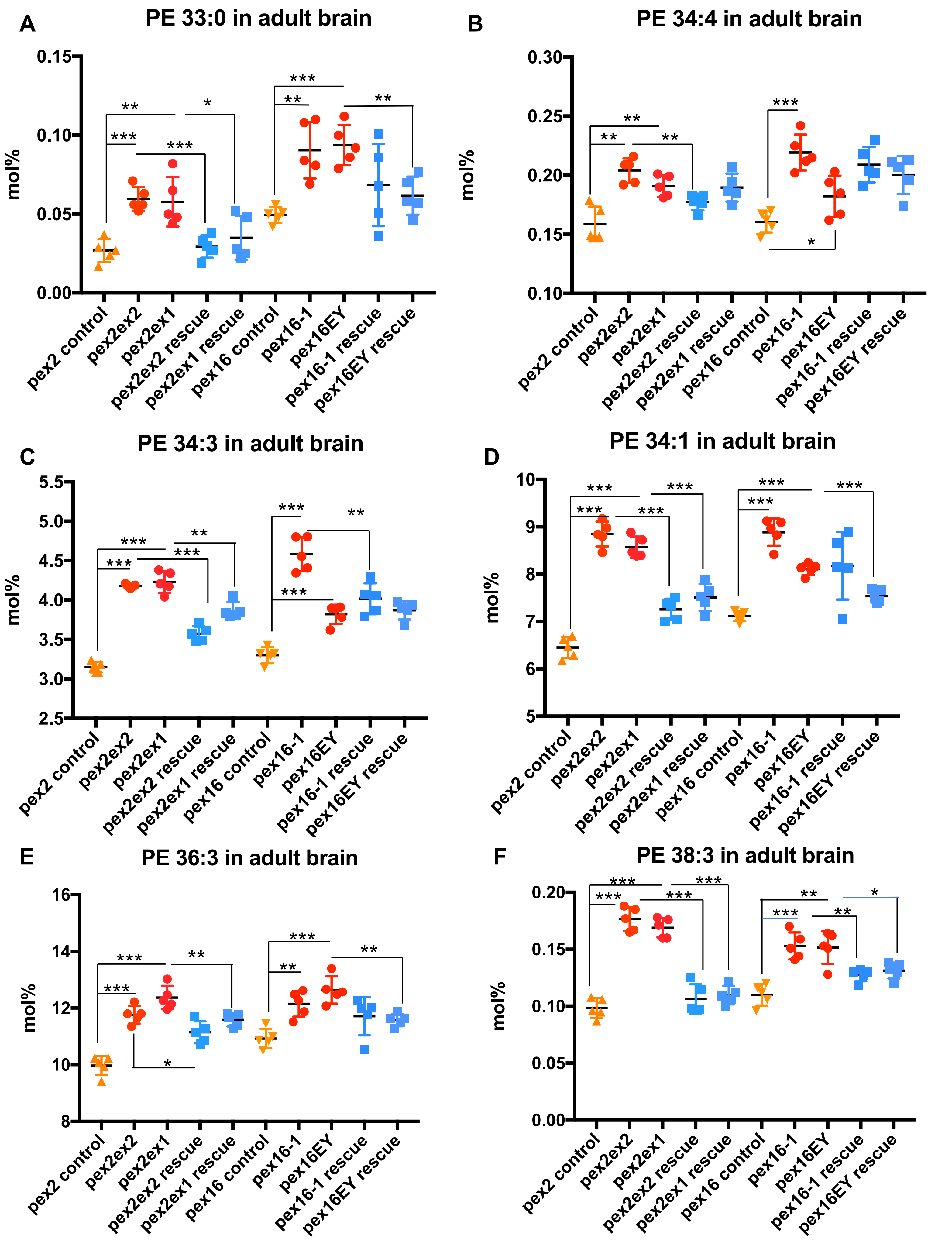

### Figure S9

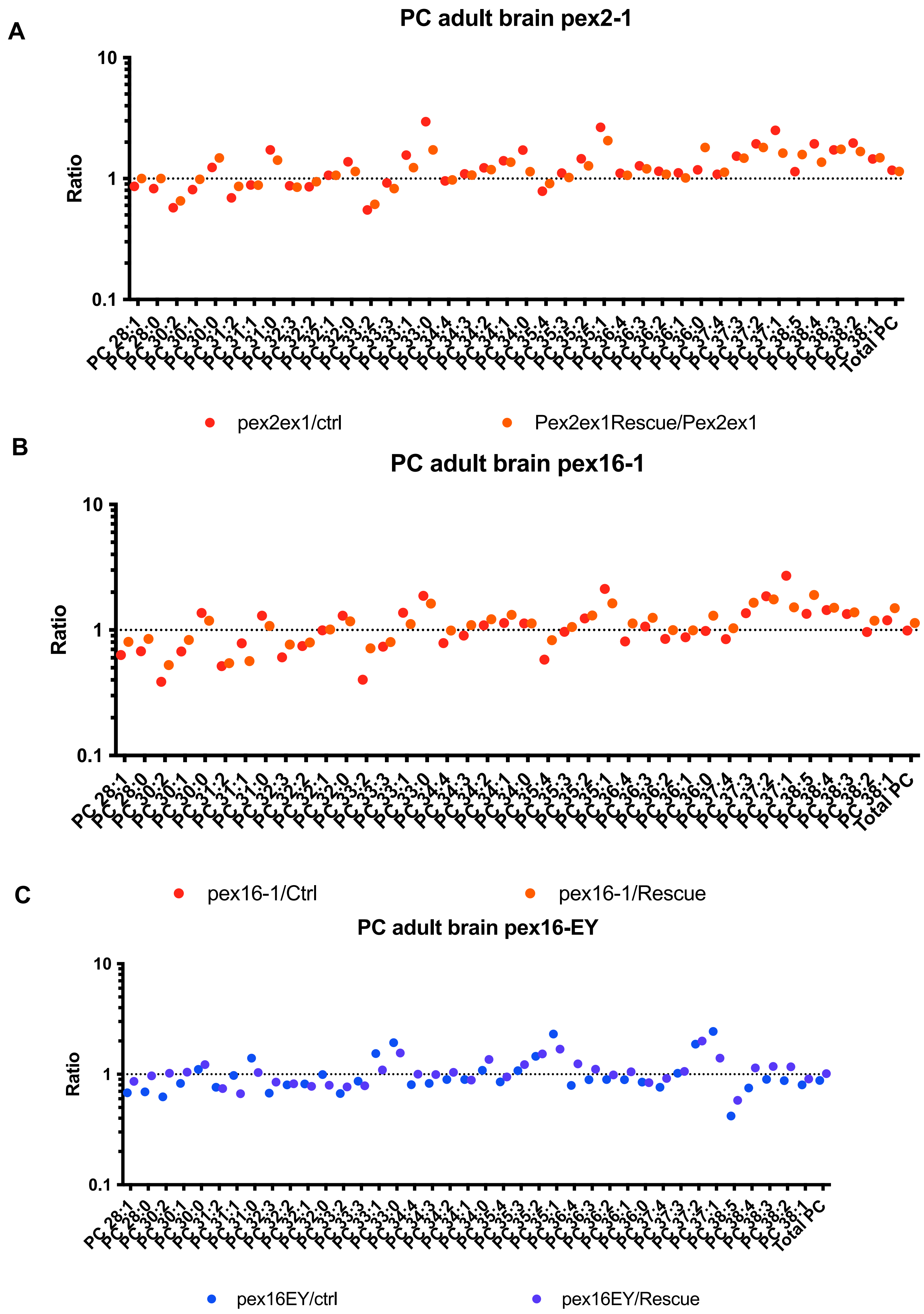

### Figure S10

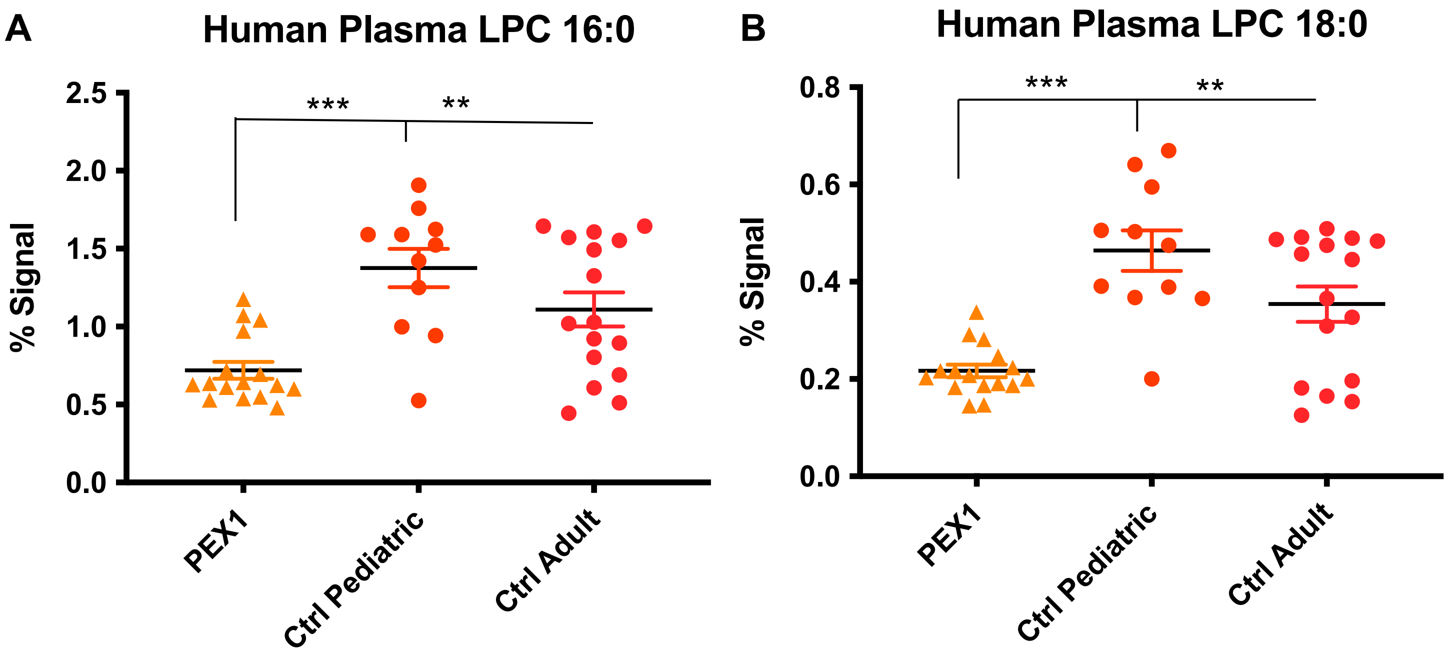

### Figure S11

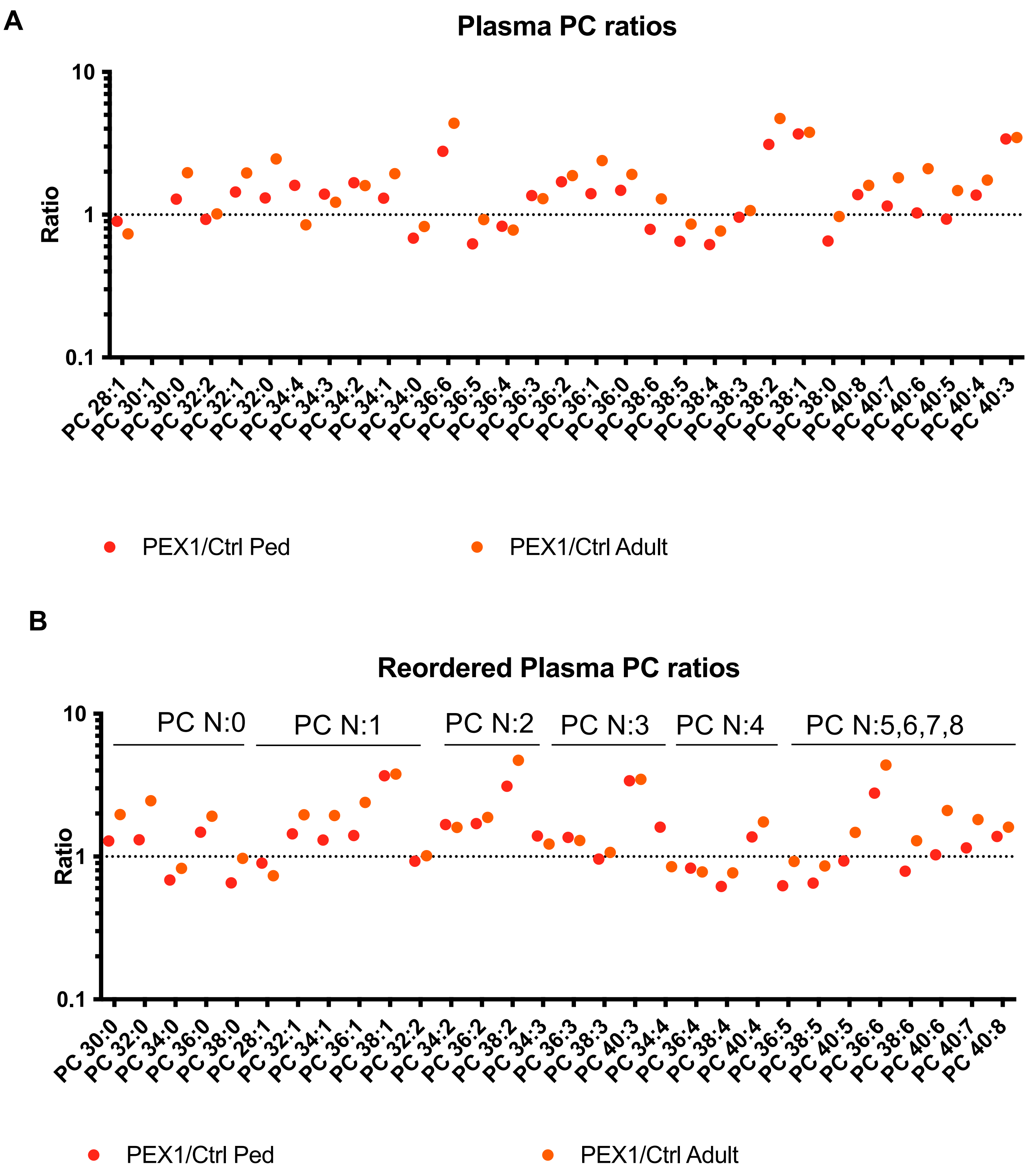
